# IL-3-driven T cell-basophil crosstalk enhances anti-tumor immunity

**DOI:** 10.1101/2024.02.01.578302

**Authors:** Jian Wei, Colleen L. Mayberry, Xiaoting Lv, Fangyan Hu, John D. Sears, Taushif Khan, Natalie A. Logan, John J. Wilson, Damien Chaussabel, Chih-Hao Chang

## Abstract

Cytotoxic T lymphocytes (CTLs) are pivotal in combating cancer, yet their efficacy is often hindered by the immunosuppressive tumor microenvironment, resulting in exhaustion. This study investigates the role of interleukin (IL)-3 in orchestrating anti-tumor immunity through CTL modulation. Intratumoral CTLs undergo a progressive decline in IL-3 production, which is correlated with impaired cytotoxic function. Augmenting IL-3, through intraperitoneal administration, IL-3-expressing melanoma cells, or IL-3-engineered CD8^+^ T cells, confers protection against tumor progression, concomitant with increased CTL activity. CTLs are critical in this therapeutic efficacy as IL-3 demonstrates no impact on tumor growth in RAG1 knockout mice or following CD8^+^ T cell-depletion. Rather than acting directly, CTL-derived IL-3 exerts its influence on basophils, synergistically amplifying anti-tumor immunity within CTLs. Introducing IL-3-activated basophils retards tumor progression, whereas basophil depletion diminishes the effectiveness of IL-3 supplementation. Furthermore, IL-3 prompts basophils to produce IL-4, which subsequently elevates IFN-γ production and viability of CTLs. Notably, the importance of basophil-derived IL-4 is evident from the absent benefits in IL-3-supplementated, IL-4 knockout tumor-bearing mice. Overall, this research unveils IL-3-mediated CTL-basophil crosstalk in regulating anti-tumor immunity and offers the prospect of harnessing IL-3 sustenance as a promising approach for optimizing and enhancing cancer immunotherapy.

**Significance Statement:** This study elucidates the critical role of IL-3 in orchestrating anti-tumor immunity, particularly within the context of CTLs and melanoma growth. It reveals a progressive decline in CTL-derived IL-3 during tumor progression, correlated with CTL exhaustion—a formidable barrier in cancer immunotherapy. Intriguingly, augmentation of IL-3, achieved through diverse means, effectively impedes tumor progression by enhancing CTL activity. This research unveils a novel mechanism: IL-3-mediated crosstalk between CTLs and IL-4-producing basophils, resulting in the rejuvenation of CTLs and amplifying their anti-tumor ability. These insights hold promise for the advancement and optimization of cancer immunotherapeutic strategies, deepening our comprehension of CTL dynamics within the tumor microenvironment, and advancing our ability to combat cancer effectively.

## INTRODUCTION

Cytotoxic T lymphocytes (CTLs) are essential in adaptive immune defense against cancers by recognizing tumor cell-specific antigens and initiating tumor cell destruction through the release of cytotoxic molecules (1, 2). The advancements in CTL research have yielded highly encouraging immunotherapeutic strategies for tackling cancer, notably encompassing immune checkpoint blockade and the adoptive transfer of tumor-reactive T cells (3–5). However, the tumor microenvironment (TME) can significantly impact the activity of tumor-infiltrating CD8^+^ T cells. Despite their initial success in fighting the disease, CTLs often progressively differentiate into an exhausted state characterized by sustained expression of inhibitory receptors and reduced production of effector cytokines (6, 7). This exhaustion impairs the anti-tumor capability of CTLs, presenting a major barrier to the success of cancer immunotherapies (8–10). The TME is composed of complex interactions among various components, such as tumor cells, immune cells, stromal cells, extracellular matrix, and numerous molecules (11). CTL exhaustion is thought to be driven by factors within tumors, including persistent antigen stimulation, co-regulatory signals, immunosuppressive cells, and cytokines (11, 12). Additionally, metabolic insufficiency imposed by nutrient deprivation and hypoxia in the TME contributes to CTL dysfunction (13–19). However, much remains unknown about the cellular and molecular mechanisms that govern CTL activity in the TME; therefore, understanding these mechanisms will offer new opportunities for the improvement of existing cancer immunotherapies and the generation of novel treatment options.

Cytokines play a critical role in regulating cell communication over short distances and shaping immune responses. Beyond mediating tumor cell death directly, CTLs also secrete cytokines with anti-tumorigenic properties, such as interferon gamma (IFN-γ), tumor necrosis factor alpha (TNF-α), and interleukin (IL)-2 (1, 2). These cytokines contribute to the effector phase of the immune response against tumors, promoting the activation of other immune cells and enhancing the anti-tumor immune response. However, CTLs frequently lose their ability to produce sufficient effector cytokines as tumor growth progresses (20). IL-3 is a pluripotent cytokine primarily produced by activated T cells (21–23). Its receptor is composed of a ligand-specific alpha subunit and a signal-transducing beta subunit, the latter of which is shared with receptors of IL-5 and granulocyte-macrophage colony stimulating factor (GM-CSF) (24, 25). IL-3 elicits the production of a variety of blood cells from the bone marrow, including granulocytes, monocytes, macrophages, eosinophils, basophils, and mast cells (26–30). Although IL-3 is dispensable for normal hematopoiesis, as evidenced by the absence of significant hematopoietic defects in IL-3-deficient mice (31), it plays an important role in immune responses during pathogen clearance and inflammation (32, 33). Introduction of the *Il3* gene into murine tumor cells has been shown to increase immunogenicity and inhibit tumor progression (34–38), thus demonstrating the potential of IL-3 to enhance anti-tumor responses. While IL-3-based anti-cancer therapies hold promise, the cellular targets and underlying mechanisms by which IL-3 exerts anti-tumor efficacy are not well defined, largely due to the broad range of leukocytes activated by IL-3. As a result, current knowledge concerning IL-3 generation by tumor-resident CTLs and its potential application in cancer therapies has been limited.

In this study, we harnessed genetically modified CTLs and tumor cells to finely modulate IL-3 levels, creating a precisely controlled experimental setting for investigating the role of IL-3 in shaping anti-tumor immunity and assessing its potential for enhancing cancer immunotherapy. Our findings revealed that CTLs within tumors progressively lose their ability to express IL-3, which is crucial for generating a potent anti-tumor response. Moreover, we identified that this deficiency is associated with increased CTL exhaustion. Supplementation of IL-3 by either intraperitoneal administration or T-cell transfer prevented tumor progression and was dependent on augmented activity of CTLs. We uncovered that IL-3 orchestrates a crosstalk between T cells and IL-4-producing basophils to enhance survival and anti-tumor function of CTLs. These results offer novel insights into the mechanism of CTL-derived IL-3 in regulating anti-tumor immunity and present new avenues for cancer immunotherapeutic approaches.

## RESULTS

### IL-3 production by T cells progressively declines during tumor growth

The B16 melanoma model was utilized to investigate the development of T-cell exhaustion during early to advanced stage tumor growth. To determine the progression of T-cell exhaustion, B16-OVA melanomas were harvested from C57BL/6J tumor-bearing mice on days 8, 13, 18, and 23 p.t.i. and tumor-resident T cells were isolated and analyzed by flow cytometry. As tumor growth progressed the expression of PD-1 and TIM-3 increased, T-bet decreased, and the production of effector cytokines (IFN-γ, TNF-α, and IL-2) decreased, indicating progressive development of T-cell exhaustion during tumor growth (**Figures S1A-S1F**). Moreover, we observed that both *in vitro* activated CD8^+^ and CD4^+^ T cells exhibited high levels of IL-3 expression, relative to their naive counterparts (**Figures S2A and S2B**). Notably, substantial IL-3 production was also evidenced in CD8^+^ and CD4^+^ T cells derived from early-stage tumors (8 days p.t.i.), contrasting with the naïve T cells isolated from the spleen. However, it is important to note that there was a progressive decline in IL-3 levels over the course of tumor development (**Figures 1A, 1B, S2C, and S2D**). Analysis of IL-3 levels and activity of CD8^+^ T cells in B16-OVA melanoma revealed that the frequencies of IL-3-producing T cells were inversely correlated with PD-1 and TIM-3 protein expression (**Figures 1C and 1D**), and were positively correlated with IFN-γ production (**Figure 1E**), suggesting that a decline in IL-3 abundance is closely linked with the progression of T-cell exhaustion in the TME. This IL-3 decline may be attributed to prolonged stimulation of T cell receptor (TCR) signaling by persistent tumor antigen and glucose deprivation in the TME, which leads to impaired anti-tumor function of CD8^+^ T cells (13, 39). To further understand this mechanism, we examined the IL-3 abundance level of *in vitro* cultured CD8^+^ T cells that were subject to extended activation with anti-CD3/CD28 antibodies and low glucose conditions, to mimic the TME (14, 40). T cells that underwent 6 days of continuous stimulation exhibited much lower IL-3 expression capability compared to those that experienced only 3 days of stimulation (**Figures 1F and S2E**). In addition, limiting glucose availability for activated T cells, through imposed low glucose culture conditions, greatly reduced IL-3 expression at both the protein and mRNA levels, suggesting an important role for glucose deprivation in T-cell IL-3 regression under persistent antigen stimulation (**Figures 1G, S2F and S2G**). Taken together, these data indicate that the decline in IL-3 abundance in exhausted T cells can be attributed to tumor-imposed environmental challenges.

**Figure 1.**
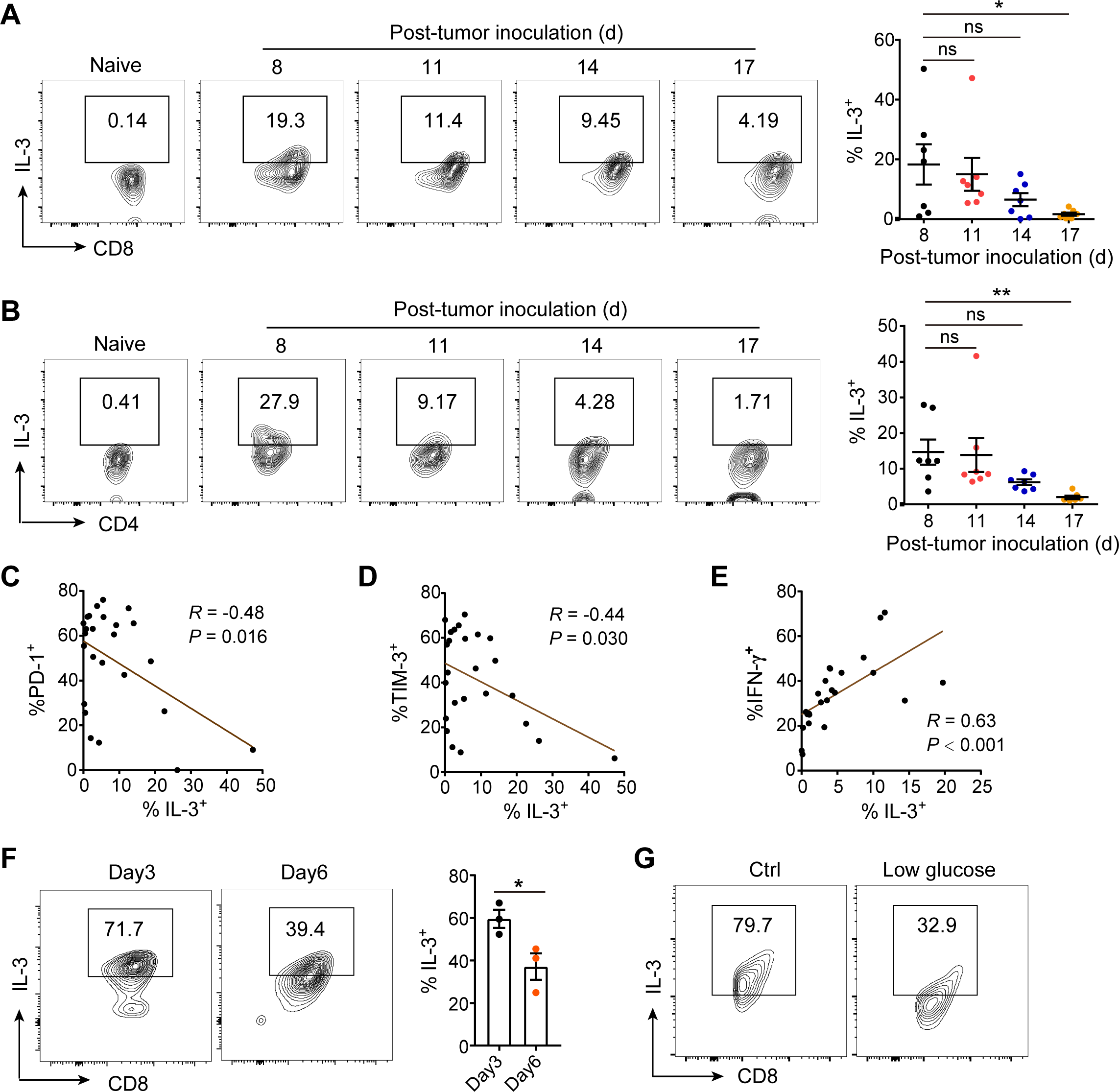
TME-imposed T cell exhaustion results in suppressed expression of IL-3. (**A**, **B**) IL-3 production by naive and CD8^+^ T cells (**A**) as well as CD4^+^ T cells (**B**) in B16-OVA melanoma at day 8, 11, 14, and 17 p.t.i was analyzed by flow cytometry. Representative flow cytometry plots (left) and percentages (right) of IL-3^+^ cells are presented (n = 7). Statistical data are presented as the mean ± SEM. (**C**) Correlation analysis between PD-1^+^ and IL-3^+^ percentages of CD8^+^ T cells in B16-OVA melanoma at day 13 p.t.i (n = 25). (**D**) Correlation analysis between TIM-3^+^ and IL-3^+^ percentages of CD8^+^ T cells in B16-OVA melanoma at day 13 p.t.i (n = 25). (**E**) Correlation analysis between IFN-γ^+^ and IL-3^+^ percentages of CD8^+^ T cells in B16-OVA melanoma at day 13 p.t.i (n = 25). (**F**) IL-3 production by day 3- and day 6-activated CD8^+^ T cells with anti-CD3/CD28 stimulation. Representative flow cytometry plots (left) and percentages (right) of IL-3^+^ cells are presented. Statistical data are presented as the mean ± SEM (n = 3). (**G**) IL-3 production by *in vitro* activated CD8^+^ T cells under glucose restriction. Representative flow cytometry plots are presented. Significance was determined by one-way ANOVA (**A**, **B**), simple linear regression (**C-E**), or Student’s *t*-test (**F**). ns, no significance; \**p* < 0.05, and \*\**p* < 0.01. See also Figure S1 and Figure S2.

### IL-3 supplementation effectively mediates tumor growth control

IL-3 expression in tumor infiltrating T cells diminishes during tumor growth (**Figures 1** and **S1**). To investigate whether supplementation of IL-3 abrogates tumor growth, we initially administered recombinant IL-3 via intraperitoneal (i.p.) injection in B16-OVA-tumor-bearing mice. Tumor growth was monitored, and at 17 days p.t.i., the mice were euthanized, and tumors were harvested and weighed (**Figure 2A**). Treatment with recombinant IL-3 resulted in a significant inhibition of tumor growth, as evidenced by smaller tumors at endpoint (**Figures 2B and 2C**). Next, to better understand the implications of IL-3 abundance in the TME, B16 melanoma cells were genetically engineered to overexpress *Il3* (B16-IL3). Subsequently, the B16-IL3 or mock control tumor cells were injected into mice, and tumor growth and survival were assessed. A significant reduction in tumor growth was measured, and an enhanced host survival rate was observed in B16-IL3 tumor-bearing mice when compared to the mock control (**Figures 2D and 2E**). Notably, even in the presence of a mixed population of tumor cells (1% B16-IL3 and 99% wildtype B16), tumor growth was restrained, indicating the robustness of IL-3 in anti-tumor activity (**Figures 2F and 2G**). Furthermore, to investigate whether IL-3 overexpression contributes to metastatic melanoma tumorigenesis, B16-IL3 or mock cells were intravenously injected into mice. Livers were harvested 20 days p.t.i. and the number of nodules was determined. Consistently, B16 cells with heightened IL-3 expression formed significantly fewer metastatic liver carcinoma nodules compared to mice injected with mock cells (**Figure 2H**), suggesting that IL-3 supplementation orchestrates a reduction in melanoma metastasis. Lastly, the influence of CTL-derived IL-3 supplementation in the TME was determined. CD8^+^ T cells with TCR designed to recognize OVA peptide residues 257-264 (OT1 cells) were modified to overexpress IL-3, and then were subsequently transferred into B16-OVA-tumor-bearing mice (**Figure 2I**). IL-3 overexpression in OT1 cells led to a substantial enhancement of tumor-restraining function, resulting in slower tumor growth and prolonged survival of the recipients compared to those receiving mock-transduced OT1 cells (**Figures 2J and 2K**). Taken together, these data suggest that IL-3 supplementation either alone or in combination with T-cell transfer, possess robust activity in control of tumor growth.

**Figure 2.**
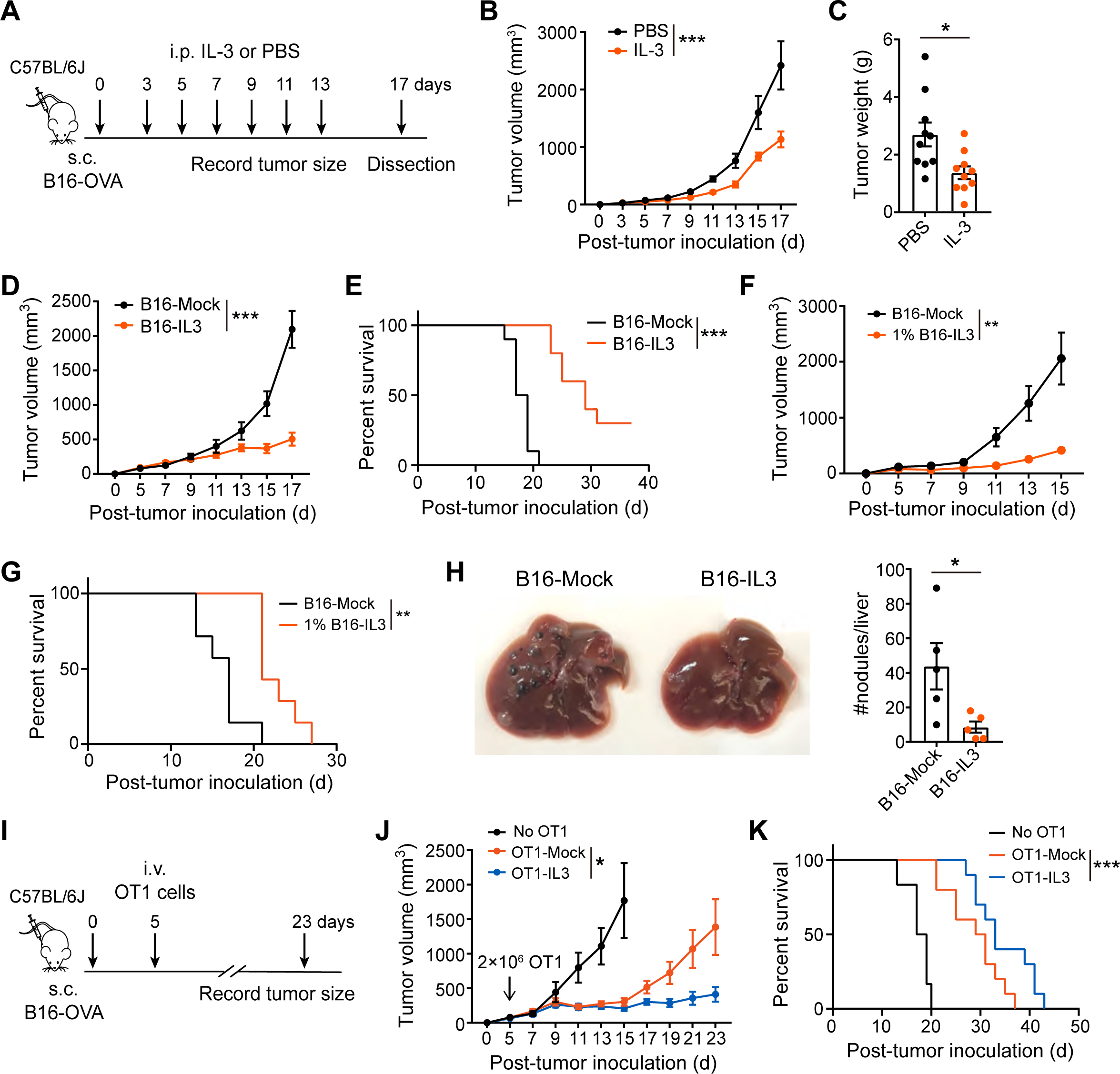
IL-3 supplementation effectively restrains tumor growth. (**A-C**) C57BL/6J mice were inoculated with B16-OVA cells by subcutaneous (s.c.) injection. The B16-OVA-melanoma-bearing mice were intraperitoneally (i.p.) administrated with IL-3 or PBS every other day from day 3 to 13 p.t.i (**A**). Tumor growth was monitored until the mice were sacrificed at day 17 p.t.i (**B**) when the tumors were isolated and weighted (**C**) (n = 10). Statistical data are presented as the mean ± SEM. (**D, E**) The growth of B16-OVA-melanoma overexpressing IL-3 (B16-IL3) or Mock (B16-Mock) (**D**) and survival curves of the tumor-bearing mice (**E**) (n = 10 mice/group). (**F, G**) Growth of B16-OVA-melanoma containing 1% B16-OVA cells overexpressing IL-3 (**F**) and survival of the tumor-bearing mice (**G**) (n = 7 mice/group). (**H**) B16-OVA cells overexpressing IL-3 or Mock were intravenously injected into mice to generate metastases. 20 days later, the mice were sacrificed, and livers were collected and examined for nodules. Representative photographs of the livers with tumor metastases (left) and graphical representation of nodule number/liver (right) are presented (n = 5). (**I-K**) OT1 cells overexpressing IL-3 or Mock were adoptively transferred by intravenous (i.v.) injection into mice with B16-OVA-melanoma at day 5 p.t.i (**I**). Tumor growth (**J**) and mice survival (**K**) were monitored (n = 6-10 mice/group). Statistical significance was determined by two-way ANOVA (**B**, **D**, **F**, and **J**), Log-rank (**E**, **G**, and **K**), or Student’s *t*-test (**C**, **H**). \**p* < 0.05, \*\**p* < 0.01, \*\*\**p* < 0.001.

### IL-3 indirectly enhances CD8^+^ T-cell anti-tumor activity

While IL-3 supplementation, either through direct injection or through T-cell mediated delivery, leads to diminished tumor growth (**Figure 2**), the underlying mechanism by which this amelioration occurs is unclear. Thus, we sought to uncover the IL-3-triggered immunological alterations that are responsible for the anti-tumor outcome. To address this, we initially assessed the requirement of T and B immune cell population contributions to the antitumor effect. We introduced 1% B16-OVA cells overexpressing IL-3 (a mixture of 1% IL-3 overexpressing and 99% wildtype B16-OVA), or solely mock control B16-OVA into B6 or Rag1 KO mice (lack mature T or B cells) and measured tumor growth over time. In wildtype (WT) mice, the growth of B16 melanomas containing 1% cells expressing IL-3 was significantly slower than that of mock control, while IL-3 expression had no impact on tumor development in the Rag1 KO mice (**Figure 3A**). This suggests that IL-3-mediated anti-tumor activity is dependent on the functionality of lymphocytes and does not exert a direct inhibitory effect. To investigate the specific contribution of CD8^+^ T cells to the anti-tumor role of IL-3, CD8^+^ T cells were depleted in mice utilizing an anti-CD8a neutralizing monoclonal antibody. We then examined the progression of tumors with or without tumor-imposed 1% IL-3 overexpression (**Figures 3B and 3C**). Depletion of CD8^+^ T cells abrogated the slower tumor growth and improved survival of 1% IL-3 B16-OVA tumor-bearing mice, highlighting the essential role of CD8^+^ T cells in IL-3-mediated tumor control (**Figures 3D and 3E**). Furthermore, we investigated the effect of recombinant IL-3 on tumor-infiltrating CD8^+^ T-cell activity (**Figure 3F**). IL-3 administration led to significant increases in the expression of CD44, T-bet, proliferation marker Ki67, IFN-γ, and TNF-α of CD8^+^ T cells, with no effect on production of cytolytic protein granzyme B (GZMB) (**Figures 3G-L**). Additionally, there was a robust reduction in PD-1 levels and a moderate decrease in TIM-3 on CD8^+^ T cells from mice that received intraperitoneal injection of recombinant IL-3 (**Figures 3M and 3N**). These data suggest that IL-3 supplementation restricted tumors by invigorating the effector activity of CD8^+^ T cells and alleviating T-cell exhaustion.

**Figure 3.**
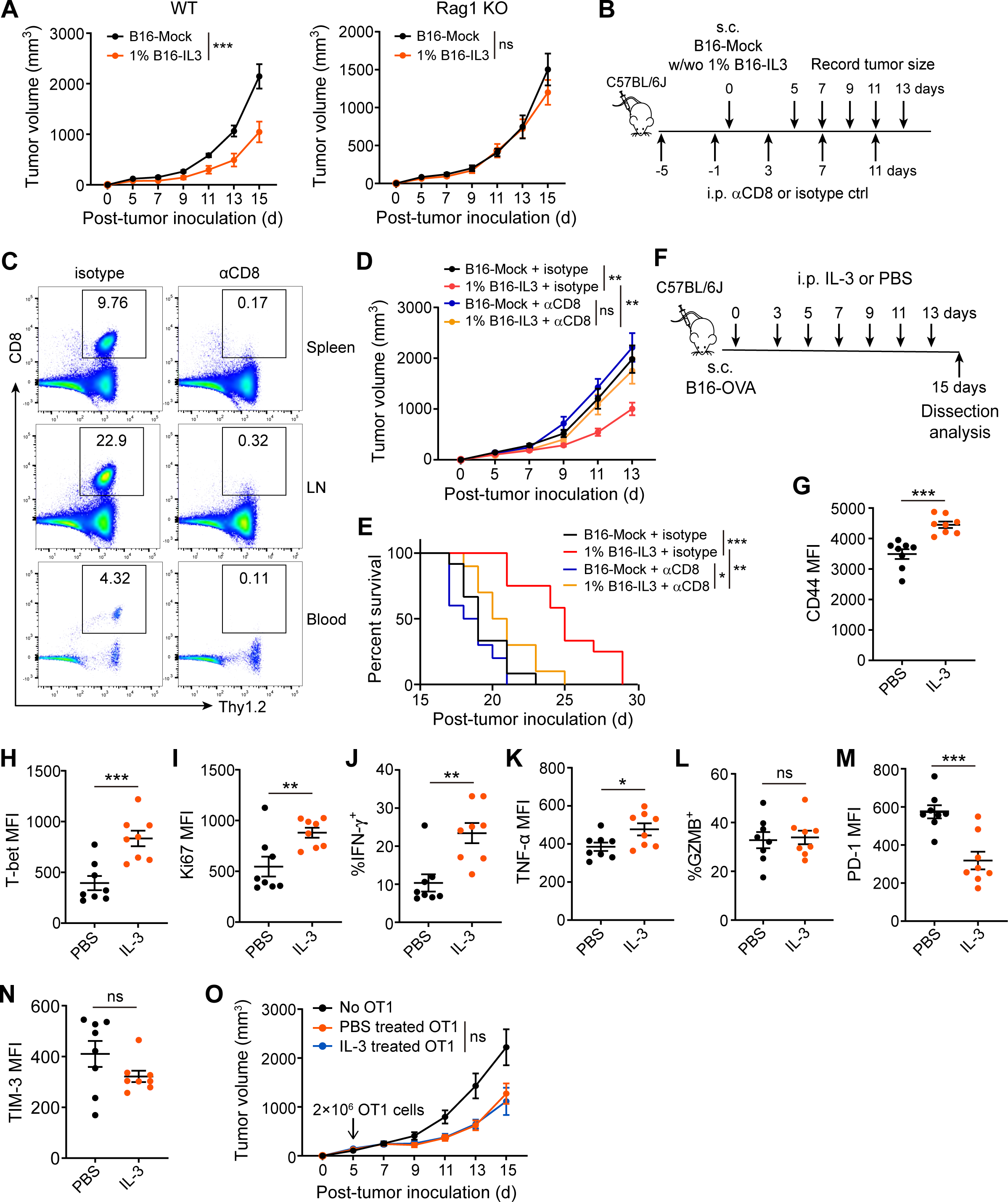
CD8^+^ CTLs are essential for IL-3 anti-tumor effects. (**A**) The growth of B16-OVA-melanoma with or without 1% B16-OVA cells overexpressing IL-3 in C57BL/6J wildtype (WT) and Rag1 KO mice (n = 7 mice/group). (**B-E**) Mice treated with isotype control IgG or αCD8 were inoculated with B16-Mock cells with or without 1% B16-IL3 cells by subcutaneous (s.c.) injection (**B**). Frequencies of CD8^+^ T cells in spleen, lymph node (LN), and blood were analyzed by flow cytometry (**C**). Tumor growth (**D**) and mouse survival (**E**) were recorded (n = 10). (**F-N**) B16-OVA-melanoma-bearing mice were intraperitoneally (i.p.) administrated with IL-3 or PBS every other day from day 3 to 13 p.t.i (**F**). Frequencies or MFI measurements of CD44 (**G**), T-bet (**H**), Ki67 (**I**), IFN-γ (**J**), TNF-α (**K**), GZMB (**L**), PD-1 (**M**), and TIM-3 (**N**) on tumor-infiltrating CD8^+^ T cells were examined at day 15 p.t.i (n = 8). (**O**) B16-OVA tumor growth following adoptive transfer of PBS or IL-3-treated OT1 cells (n = 6-7 mice/group). Statistical significance was determined by two-way ANOVA (**A**, **D**, and **O**), Log-rank (**E**), or Student’s *t*-test (**G-N**). ns, no significance; \**p* < 0.05, \*\**p* < 0.01, \*\*\**p* < 0.001. See also Figure S3.

To gain deeper insight into the potential direct effect of IL-3, we subjected activated CD8^+^ T cells to recombinant IL-3 treatment and evaluated their metabolism, effector function, and viability. CD8^+^ T cells in the presence or absence of IL-3 exhibited similar metabolic patterns, including glucose uptake, glycolysis rates, oxidative phosphorylation (OXPHOS) rates, and glycolysis/OXPHOS ratios (**Figures S3A-S3D**). Additionally, similar states of activation were observed, as profiled by markers CD44, PD-1, TIM-3, CD25, T-bet, and Ki67 (**Figures S3E-S3J**), comparable production of cytokines (including TNF-α, IL-2, IFN-γ, and GZMB) (**Figures S3K-S3N**), and similar viability (**Figure S3O**). Accordingly, IL-3 pre-treated T cells exerted an anti-tumor capability equivalent to those treated with PBS (**Figure 3O**). Collectively, these results suggest that IL-3 did not directly influence T-cell activity, but rather enhanced the anti-tumor immunity of CD8^+^ T cells through an intermediary mechanism, signifying an indirect impact.

### IL-3-mediated crosstalk between T-cells and non-B/T cells enhances anti-tumor capacity of CD8^+^ T cells

Our findings suggest that IL-3-mediated enhancement of CD8^+^ T cell anti-tumor activity may involve the engagement of intermediary cell populations (**Figure 3**). To evaluate the influence of accessory cells on the activation of CD8^+^ T cells, we compared the activation of purified CD8^+^ T cells (pCD8) to that of CD8^+^ T cells in the presence of bulk splenocytes (bCD8). We found that, compared to pCD8 cells, bCD8 cells displayed glycolysis-dominant metabolism, characterized by increased glucose uptake, higher glycolytic rate, lower OXPHOS rate, and elevated ECAR/OCR ratio (**Figures S4A-S4D**). In addition, the *in vitro* cultured bCD8 cells displayed enhanced IFN-γ production (**Figure S4E**) and improved survival post activation (**Figure S4F**). Moreover, when adoptively transferred into B16-OVA-bearing mice, the frequency of tumor-infiltrating bCD8 OT1 cells was significantly higher than that of pCD8 OT1 cells at 4 days post-transfer (**Figures S4G and S4H**). Consequently, relative to pCD8 OT1 cells, bCD8 OT1 cells possessed greatly superior cytotoxicity against B16-OVA tumor cells in both *in vitro* and *in vivo* conditions (**Figures S4I-S4K**). Overall, these results demonstrate that the presence of accessory cells positively impacts the protective anti-tumor immunity of CD8^+^ T cells.

To further investigate the underlying mechanism of this positive impact, we employed a permeable transwell system to determine whether the modulation of T cells by bulk accessory cells was mediated by cell-cell contact or via cellular secreted factors. Our findings revealed that CD8^+^ T cells cultured in transwell with bulk splenocytes (pCD8 (TW)) exhibited a similar phenotype to bCD8 cells, as evidenced by increased IFN-γ production and improved viability relative to pCD8 cells cultured alone (**Figures 4A and 4B**). This indicates that soluble factors released by other cell populations mediate the enhancement of CD8^+^ T-cell effector function. Despite this fact, conditioned medium from culture of activated CD8^+^ T cells (CD8^+^ T-sup) or Rag1 KO mouse splenocytes (Rag-sup) did not have the ability to raise IFN-γ production and viability of CD8^+^ T cells (**Figures 4C and 4D**). However, when Rag1 KO mouse splenocytes were previously incubated with conditioned medium of CD8^+^ T cells, their supernatant acquired the ability to facilitate CD8^+^ T-cell IFN-γ production and survival relative to the medium control (**Figures 4C and 4D**). This suggests that T cell-derived soluble effectors enabled certain non-T/B cell subsets to improve the effector function of CTLs. We hypothesized that IL-3, a soluble factor primarily produced by activated T cells, may play a role in mediating this interaction. To test this hypothesis, we first determined whether the role of T-cell conditioned medium could be functionally substituted by IL-3. The results showed that, although IL-3 had no effect when directly applied to CD8^+^ T cells, conditioned medium of IL-3-treated Rag1 KO mouse splenocytes improved IFN-γ production and viability of CTLs (**Figures 4E and 4F**). Despite sharing the same beta subunit with the IL-3 receptor, neither IL-5 nor GM-CSF were able to recapitulate the effect observed with IL-3 in augmenting IFN-γ production and survival of T cells in splenocytes from Rag1 KO mice (**Figures S4L and S4M**). To further investigate this, we assessed the necessity of IL-3 in the crosstalk of T-cell and other cells utilizing αIL-3 neutralizing antibody. The inclusion of neutralizing IL-3 antibodies in the conditioned medium of activated T cells greatly diminished the beneficial effects on both IFN-γ production and survival of CD8^+^ T cells (**Figures 4G-4I**), heavily suggesting that IL-3 produced by CTLs activates other cell subsets that, in turn, augment the T-cell effector function. Taken together, these data firmly established an IL-3-mediated crosstalk between T-cell and non-B/T cells, ultimately contributing to the enhancement of anti-tumor capability of CTLs.

**Figure 4.**
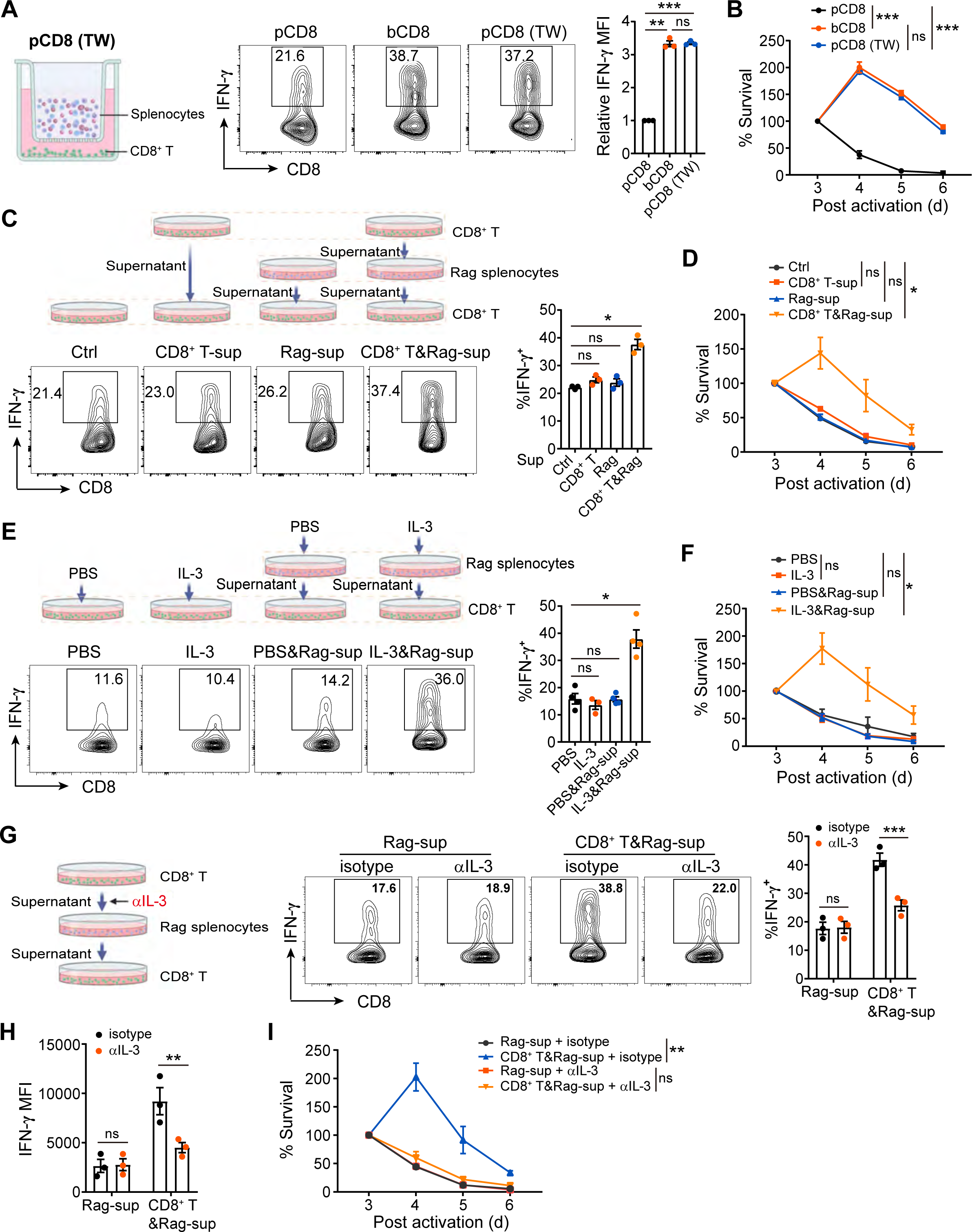
T-cell-derived IL-3 indirectly augments effector function of CD8^+^ T cells via stimulating non-B/T cells. (**A, B**) Purified CD8^+^ T cells were activated and cultured with bulk splenocytes in transwell plates (pCD8(TW)) as indicated. IFN-γ production (**A**) and survival (**B**) of the cells were evaluated (n = 3). (**C, D**) Purified CD8^+^ T cells were activated and cultured with conditioned supernatant from culture of CD8^+^ T cells or Rag1 KO mouse splenocytes as indicated (**C**). IFN-γ production (**C**) and survival (**D**) of the cells were evaluated (n = 3). (**E, F**) Purified CD8^+^ T cells were activated and cultured with PBS, IL-3, or conditioned supernatant from Rag1 KO mouse splenocytes stimulated with PBS or IL-3 (**E**). IFN-γ production (**E**) and survival (**F**) of the cells were evaluated (n = 3). (**G-I**) Rag1 KO mouse splenocytes were cultured with CD8^+^ T-cell supernatant in the presence of isotype control IgG or αIL-3 for 2 days. Then the conditioned medium was used for the culture of purified CD8^+^ T cells (**G**). Representative flow cytometry plots and percentage of IFN-γ^+^ cells (**G**), IFN-γ MFI (**H**), and survival (**I**) of the CD8^+^ T cells were evaluated. Statistical significance was determined by one-way ANOVA (**A**, **C**, and **E**), two-way ANOVA (**B**, **D**, **F**, and **I**), or Student’s *t*-test (**G**, **H**). ns, no significance; \**p* < 0.05, \*\**p* < 0.01, \*\*\**p* < 0.001. See also Figure S4.

### IL-3-activated basophils show potency in restricting tumor growth and enhancing anti-tumor immunity

In order to identify the cell target of IL-3, we isolated various splenic cell populations by flow cytometry from Rag1 KO mice and evaluated their impact on CD8^+^ T cells following recombinant IL-3 treatment (**Figure S5A**). The purified cell subsets analyzed included CD45^-^ cells, natural killer cells (NK), neutrophils, dendritic cells (DC), macrophages, monocytes, and CD45^+^ lineage marker negative (Lin^-^) cells. Of these cell subsets, only the conditioned medium of IL-3-treated CD45^+^ Lin^-^ cells displayed the ability to enhance IFN-γ production of CD8^+^ T cells (**Figure S5B**). Similarly, CD45^+^ Lin^-^ cells isolated from tumors also enhanced IFN-γ production and viability of CD8^+^ T cells in response to IL-3 treatment, through culture of CD8^+^ T cells in CD45^+^ Lin^-^ conditioned medium (**Figures 5A and 5B**), indicating the presence of specific cells responsive to IL-3 in the TME.

**Figure 5.**
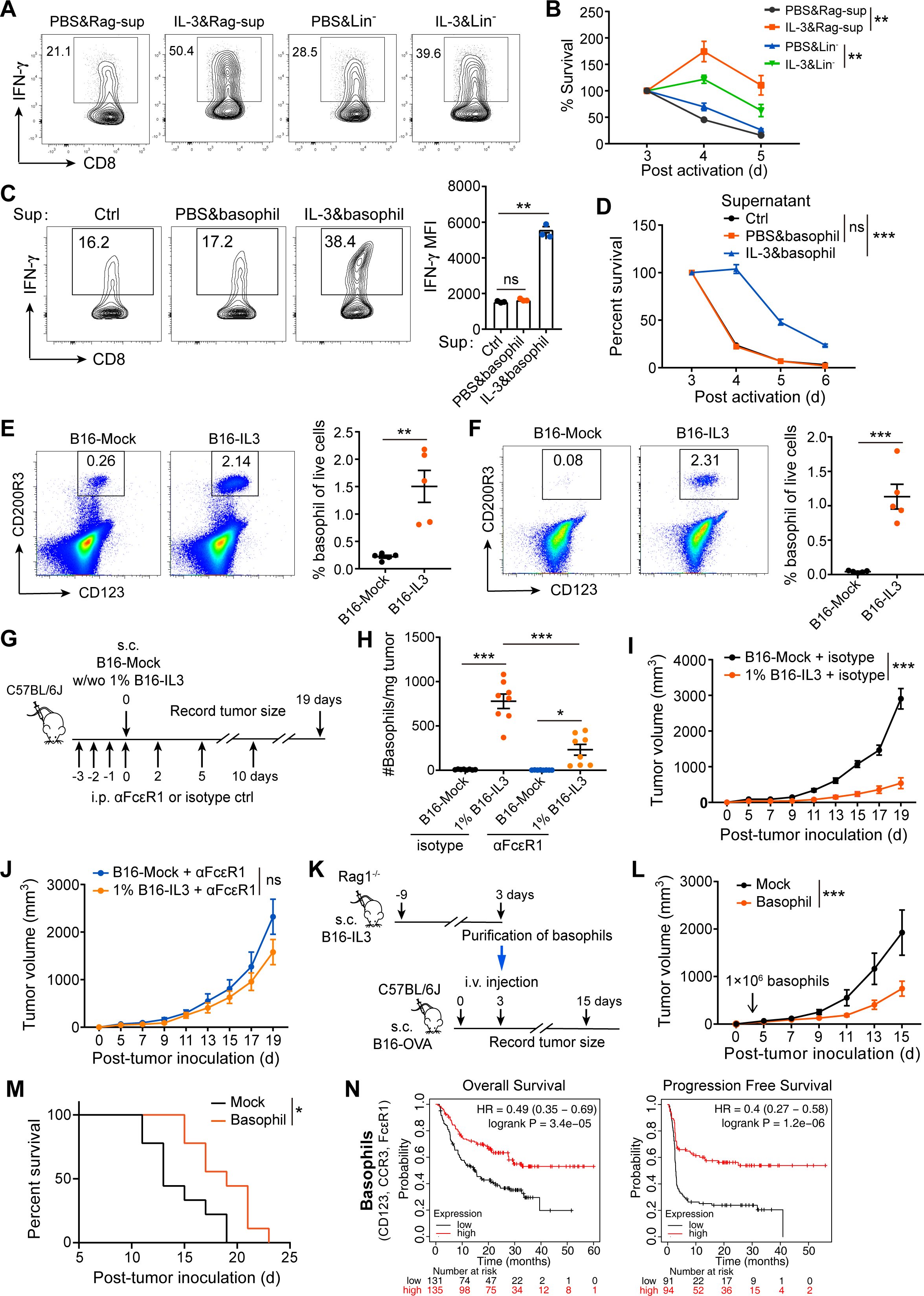
IL-3 stimulated basophils strengthen effector function of CD8^+^ T cells. (**A, B**) CD45^+^Lin^-^ cells isolated from B16 melanomas were cultured with PBS or IL-3 and the conditioned supernatant was taken for CD8^+^ T cells culture. IFN-γ production (**A**) and survival (**B**) of the CD8^+^ T cells was examined. (n = 3) (**C, D**) CD8^+^ T cells were cultured with conditioned supernatant from basophils stimulated with PBS or IL-3. IFN-γ production (**C**) and survival (**D**) of the CD8^+^ T cells were evaluated (n = 3). (**E, F**) Basophils in the spleen (**E**) and tumor (**F**) of mice with B16-Mock or B16-IL3 melanomas were evaluated at day 14 p.t.i. Representative flow cytometry plots (left) and percentage (right) of basophils are presented (n = 5). (**G-J**) Mice treated with isotype control IgG or αFcεR1 were inoculated with B16-Mock cells with or without 1% B16-IL3 cells by subcutaneous (s.c.) injection (**G**). Basophils in the tumors (**H**) and tumor growth (**I** and **J**) were monitored (n = 7-8). (**K-M**) Basophils isolated from Rag1 KO mice with B16-IL3-melanomas were adoptively transferred into B16-OVA-melanoma-bearing C57BL/6J mice at day 3 p.t.i (**K**). Tumor growth (**L**) and mice survival (**M**) were recorded (n = 9 mice/group). (**N**) Associations of tumor-resident basophils with outcomes of melanoma patients receiving no immune checkpoint therapy. Groups of patients were divided based on combined gene expression levels of selected basophil markers (CD123, CCR3, and FcεR1) in tumor, into top tertile (T3) and lower tertile (T1), while patients in the middle tertile (T2) were excluded. Patients’ overall survival and progression free survival were analyzed up to 60 months. Statistical significance was determined by two-way ANOVA (**B**, **D**, **I**, **J**, and **L**), one-way ANOVA (**C**, **H**), Student’s *t*-test (**E**, **F**), or Log-rank test (**M**). ns, no significance; \**p* < 0.05, \*\**p* < 0.01, and \*\*\**p* < 0.001. See also Figure S5

Basophils, a subset of CD45^+^ Lin^-^ leukocytes, express high levels of IL-3 receptor (41, 42), and IL-3 has been shown to facilitate their development and proliferation (32, 43). To investigate a potential role of basophils in the IL-3-mediated regulation of T cells, sort-purified basophils (CD45^+^ FcεR1^+^ CD200R3^+^ CD123^+^ c-Kit^-^, **Figure S5C**), were treated with IL-3, and cellular culture medium was subsequently collected. CD8^+^ T cells were cultured in that conditioned medium and the levels of IFN-γ expression and the cellular viability of T cells were measured. Remarkably, following culture in the IL-3-treated basophil media, IFN-γ production and viability of CD8^+^ T cells were significantly increased (**Figure 5C and 5D**). Additionally, IL-3 supplementation through tumor-imposed IL-3 expression (1% IL-3 B16-OVA) significantly expanded the basophil population in the spleen and tumor of mice, when compared with wildtype B16-OVA (**Figures 5E and 5F**). Moreover, injection of FcεR1 neutralizing antibody, resulted in the inhibition of basophil expansion (**Figures 5G and 5H**), and also reduced the anti-tumor efficacy of IL-3 (**Figures 5I and 5J**). These results strongly suggest that basophils are the effector cells that respond to IL-3 and subsequently enhance anti-tumor immunity. Furthermore, basophils were isolated from B16-IL3-melanoma-bearing Rag1 KO mice and subsequently introduced into B16-melanoma-bearing wildtype mice to evaluate the effect on tumor growth (**Figure 5K**). The adoptive transfer of these basophils effectively restricted tumor growth and significantly prolonged host survival (**Figures 5L and 5M**), highlighting the potential of IL-3-exposed basophils to curb tumor expansion. Furthermore, we evaluated the potential impact of tumor-resident basophils on patient survival outcomes in human melanoma using the basophil marker genes CD123, CCR3, and FcεR1 (44, 45). Higher expression of the basophil marker genes within tumor is associated with significantly enhanced overall survival and progression free survival among melanoma patients who did not undergo any immune checkpoint therapies (**Figure 5N**). Taken together, these findings suggest that basophils activated by IL-3 may play a crucial role in enhancing anti-tumor immunity through increased IFN-γ production and viability of CD8^+^ T cells, offering a promising approach for cancer immunotherapy.

### IL-3-induced IL-4 release from basophils promotes CD8^+^ T cell effector function and anti-tumor immunity

Our findings demonstrate that the use of conditioned medium of IL-3-treated basophils led to an enhancement in T-cell effector functions, suggesting that this effect is mediated by the release of a soluble factor (**Figure 5**). To investigate the molecular mechanism underlying this effect, we performed RNA-seq analysis of activated CD8^+^ T cells after exposure to the conditioned supernatant derived from IL-3 or PBS-exposed basophils. RNA-seq analysis indicated that a number of genes were differentially expressed in activated CD8^+^ T cells after exposure to conditioned supernatant from IL-3 or PBS-activated basophils, with some genes upregulated and others downregulated (**Figure S6A**). KEGG analysis and Gene Set Enrichment Analysis (GSEA) revealed that the primary gene expression target was the cytokine-cytokine receptor interaction (**Figures S6B and 6A**). In addition, genes involved in inflammatory response, T-cell activation, and T-cell proliferation were significantly enriched in CD8^+^ T cells exposed to supernatant from IL-3-activated basophils (**Figures 6B, S6C, and S6D**). Using Ingenuity Pathways Analysis to perform Upstream Regulator Analysis, we uncovered that the IL-4 signaling cascade (*p*= 2.73E-33, bias-corrected absolute Z score= 4.108) emerges as the foremost soluble factor exerting the significant influence on the gene expression profile of CD8^+^ T cells activated in IL-3-exposed basophil-supernatant (**Table 1**). The analysis identified 76 transcripts that were differentially regulated by IL-4 in T cells treated with IL-3-exposed basophil supernatant, with 40 decreased transcripts (green) and 36 increased transcripts (red) in response to IL-4 (**Table S1**). Of the 76 altered transcripts, 48 showed directional regulation consistent with the action of the IL-4 signaling pathway, while 15 were directionally inconsistent with regulation (yellow) and 13 displayed unpredicted regulation (gray). In line with activation of the identified IL-4 signaling cascade, genes engaged in JAK-STAT signaling, PI3K-AKT signaling, tyrosine phosphorylation of STAT protein, and intracellular signaling transduction were also enriched in T cells treated with IL-3-exposed basophil supernatant (**Figures 6C, 6D, S6E, and S6F**). These data suggest that IL-3-treated basophil-conditioned medium enhances T-cell functions potentially in response to IL-4 stimulation, resulting in significant alterations in gene expression related to cytokine signaling, inflammation, and T-cell activation.

**Figure 6.**
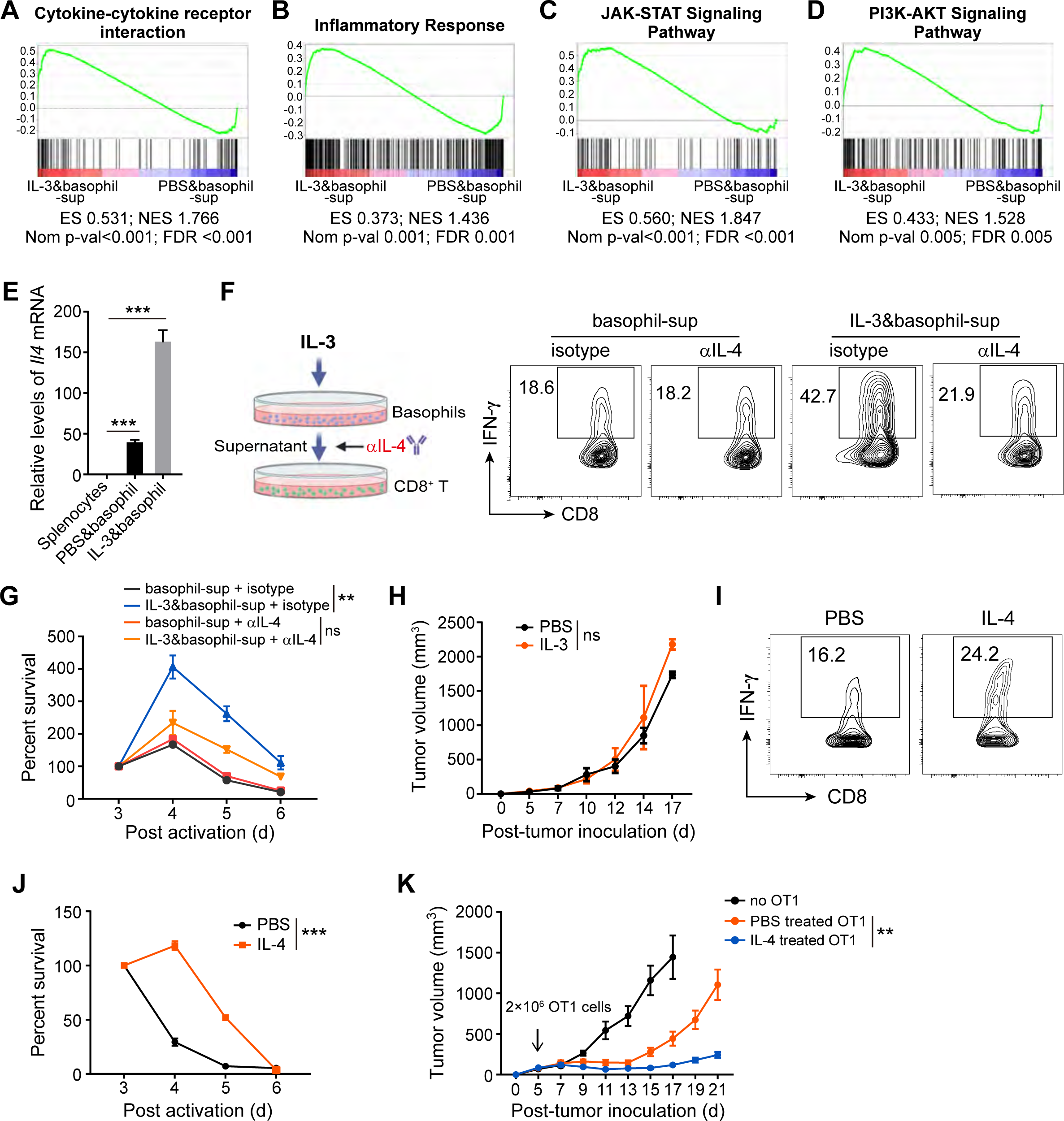
IL-3 enhances basophile IL-4 production to improve anti-tumor activity of CD8^+^ T cells. (**A-D**) GSEA plots of activated CD8^+^ T cells following incubation with conditioned supernatant from culture of basophils with IL-3 *versus* PBS (n=3 mice per group). Enrichment of gene sets in cytokine-cytokine receptor interaction (**A**), inflammatory response (**B**), JAK-STAT signaling pathway (**C**), and PI3K-AKT signaling pathway (**D**). (**E**) Relative mRNA levels of *Il4* in bulk splenocytes and basophils treated with PBS or IL-3 (n = 3). (**F, G**) CD8^+^ T cells were incubated with conditioned supernatant from IL-3 treated basophils in the presence of isotype IgG or αIL-4 (**F**). IFN-γ production (**F**) and survival (**G**) of the CD8^+^ T cells were evaluated (n = 3). (**H**) *Il4* knockout B16-OVA tumor-bearing mice were intraperitoneally (i.p.) administrated with IL-3 or PBS every other day from day 3 to 13 p.t.i and tumor growth was monitored (n = 9). (**I, J**) Purified CD8^+^ T cells were activated with αCD3/CD28 in the presence of PBS or IL-4. IFN-γ production (**I**) and survival (**J**) of the cells following activation were evaluated (n = 3). (**K**) B16-OVA tumor growth following adoptive transfer of PBS or IL-4 treated OT1 cells (n = 7 mice/group). Statistical significance was determined by one-way ANOVA (**E**), or two-way ANOVA (**G**, **H**, **J, and K**). ns, no significance; \*\**p* < 0.01, and \*\*\**p* < 0.001. See also Figure S6.

**Table 1.** Summary of identified cytokine-mediated upstream regulators.

| **Upstream Regulator\*** | **Predicted Action Status** | **Z-Score (Overlap *p*-Value)<sup>#</sup>** | **Numbers of Gene Targets** |
| --- | --- | --- | --- |
| IL-4 | Activated | 4.108 (2.73E-33) | 76 |
| IL1A | Activated | 2.732 (2.33E-06) | 14 |
| TNFSF11 | Activated | 2.578 (6.32E-09) | 18 |

\*Cytokines were identified as matching the activity of IL-3-exposed basophil supernatant using IPA upstream regulator analysis, with predicted action status and a z-score greater than 2 indicating a significant positive correlation.
<sup>#</sup>The activation z-score serves as a statistical measure and a directional indicator (positive/upregulated) while the overlap *p*-value serves as an additional measure of statistical certainty pertaining to the interrelated nature of the molecules forming each network.

Indeed, IL-3 exposure strongly augmented *Il4* gene expression in basophils by more than 3-fold when compared to the PBS-treated control (**Figure 6E**). Blockade of IL-4, via a neutralizing antibody, largely diminished the positive effects of IL-3 and basophil-supernatant in IFN-γ production and viability of CD8^+^ T cells (**Figures 6F and 6G**), highlighting the importance of IL-4 for the T cell-basophil crosstalk. This is further reinforced by data demonstrating that injection of recombinant IL-3 in *Il4* knockout B16-OVA tumor-bearing mice exerts negligible influence on tumor growth (**Figure 6H**). To better understand the effect of IL-4 on CD8^+^ T-cell activation, activated CD8^+^ T cells were cultured in the presence of IL-4, for metabolic and functional analysis. The direct exposure of activated CD8^+^ T cells to IL-4 remodeled T-cell metabolism by increasing glucose uptake, boosting glycolytic rate, reducing OXPHOS, and enhancing the ECAR/OCR ratio (**Figures S6G-S6J**). Furthermore, IL-4 treatment improved both IFN-γ production and survival of CD8^+^ T cells (**Figures 6I and 6J**). The IL-4 treatment also upregulated CD8^+^ T cell expression of CD44, while having minimal effects on the levels of PD-1, TIM-3, CD25, T-bet, Ki67, TNF-α, IL-2, and GZMB (**Figures S6K-S6S**). Importantly, adoptive transfer of tumor-reactive OT1 cells that were pre-treated with IL-4 resulted in superior anti-tumor activity against B16-OVA melanoma (**Figure 6K**), as determined by significantly reduced tumor volume. In summary, these findings demonstrate that increased production of IL-4 by IL-3-exposed basophils enhances CD8^+^ CTL-mediated anti-tumor immunity.

## DISCUSSION

Cytokines that facilitate interactions among effector cells play a crucial role in the immune response to cancer. Our study uncovers a progressive decline in IL-3 expression by CTLs during the progression of CTL exhaustion, which is attributed to persistent stimulation and/or glucose competition in the TME. Supplementation of IL-3 orchestrates the anti-tumor immunity of CTLs by enhancing crosstalk between T cells and basophils, facilitated by basophil-derived IL-4. These findings provide a cellular and molecular mechanism for anti-tumor activity of IL-3 and offer new avenues for improved cancer immunotherapy (**Figure 7**).

**Figure 7.**
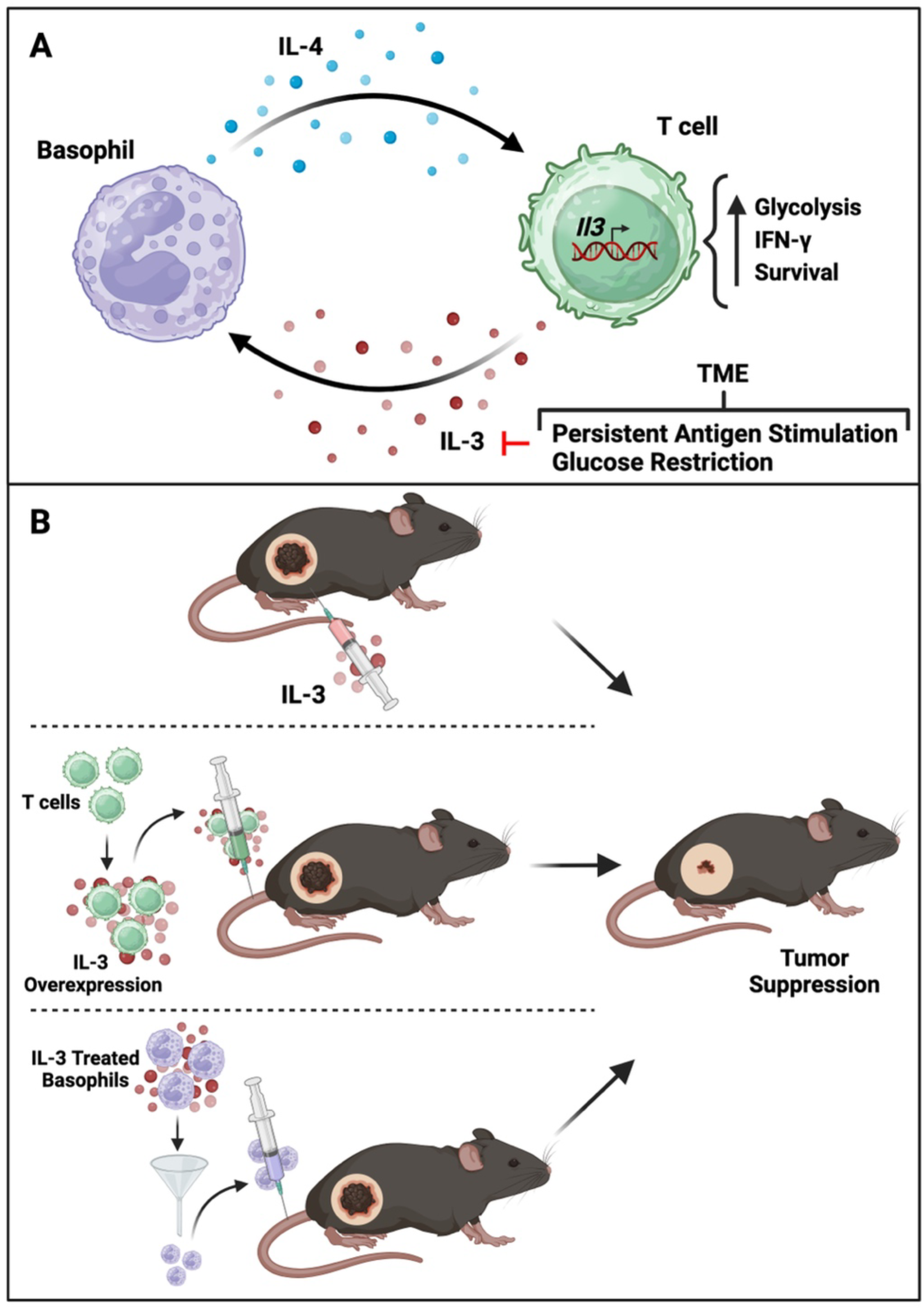
Enhanced T cell-basophil crosstalk by IL-3 supplementation improves anti-tumor immunity. Persistent antigen stimulation and glucose limitation within the TME dampen IL-3 production by CTLs over the course of tumor growth. (**A**) T cell-derived IL-3 stimulates basophils to release more IL-4, which in turn directly acts on CTLs to enhance glycolysis, IFN-γ production, and T cell viability, leading to enhanced anti-tumor immunity. IL-3-mediated T cell-basophil crosstalk sheds light on opportunities for development of improved cancer immunotherapy. (**B**) Administration of IL-3 intraperitoneal (i.p.), adoptive transfer of CD8^+^ T cells overexpressing IL-3, and transfer of IL-3-activated basophils possess the capacity to restrain tumor progression.

Previous studies have suggested that introducing the *Il3* gene into tumor cells can exert a restraining influence on tumor growth (34, 35, 37, 46). However, the precise mechanism underlying this effect remained elusive. In this study, we demonstrate that both intraperitoneal administration of recombinant IL-3 and IL-3 delivery through adoptive T-cell transfer exert strong anti-tumor activity. Critically, we found that it was the unique alpha subunit (CD123) of the IL-3 receptor that plays a crucial role in mediating the specific biological effect, as IL-5 and GM-CS, both of which share a beta subunit with IL-3, had no effect on CTL augmentation. Consistent with prior reports highlighting IL-3’s ability to enhance CTL responses in lymphoma and lung carcinoma (34–36, 47), our data demonstrate that CTLs are required for the anti-melanoma efficacy of IL-3. Furthermore, our study delves into the broader context of IL-3’s effects on the immune system; thereby revealing that direct stimulation of CTLs with IL-3 does not affect their activity and suggesting the involvement of other effector cells that respond directly to IL-3. Administration of IL-3 has been linked to the differentiation and proliferation of hematopoietic progenitors, which results in the production of multipotential cells (26–29). Additionally, IL-3-transfected lung carcinoma has been shown to enhance the development of tumor-specific CTLs by stimulating a population of host macrophage-like cells that process and present tumor antigens to T cells (35). Crucially, our findings offer significant insights into the anti-tumor efficacy of CTL-derived IL-3 and suggest that basophil expansion and cytokine secretion induced by IL-3 are critical to this effect. While IL-3 may not be essential for the steady-state generation of basophils, our research highlights its ability to activate basophils and prolong their survival, particularly at exceptionally low concentrations under certain pathological conditions (31–33, 48). Given that activated T cells are the primary source of IL-3 (21), it is reasonable to propose that, when these T cells are primed by tumor antigens, they undergo rapid proliferation and produce large quantities of IL-3. This, in turn, triggers basophil activation, thereby reinforcing a strong anti-tumor immune response.

Basophils, derived from granulocyte-monocyte progenitors in the bone marrow, constitute a relatively small fraction of the overall leukocyte population in the bloodstream (49, 50), but have been found within the TME of human lung adenocarcinoma and pancreatic cancers (51–53). However, their precise role in cancer immunity has remained enigmatic, partly due to their limited lifespan and scarce presence. Existing literature presents conflicting evidence regarding basophil function with studies suggesting involvement in melanoma rejection; conversely, these cells have been implicated in promoting tumorigenesis in pancreatic cancer (54, 55). These findings suggest that basophil involvement in cancer immunity could be tumor-specific in nature. In our study, we made the significant observation of a melanoma-associated basophil population which was greatly expanded by IL-3 treatment. Our data provide compelling support for the proposition that basophils play a pivotal role in modulating the anti-tumor response in melanoma. We further demonstrate that basophil depletion substantially diminished IL-3-initiated anti-tumor activity. Moreover, through the adoptive transfer of IL-3-activated basophils, we observed a significant deceleration in the progression of mouse melanoma, strongly implicating an anti-tumorigenic role of these basophils. Previous research has proposed multiple mechanisms to explain how basophils may exert anti-tumorigenic effects. These mechanisms encompass the release of cytotoxic molecules, such as granzyme B and TNF-α, which have the potential to induce tumor-cell death (56, 57). Additionally, some studies suggest that basophils may function as antigen presenting cells to initiate T-cell responses (58–60). Still other studies postulate that tumor-infiltrating basophils may aid in the recruitment of CD8^+^ T cells through a chemokines-dependent mechanism (54). Most critically, our data underscore the significant contribution of the IL-4 cytokine released by basophils in enhancing CTL anti-tumor activity. Together, these findings not only advance our comprehension of the intricate interplay between basophils and anti-tumor immunity, but also highlights their potential as promising therapeutics in cancer treatment.

Our findings, in conjunction with previous research (61, 62), highlight the significant role of basophils as producers of IL-4 and their profound impact on anti-tumor immunity. It is widely recognized that basophils are constitutive producers of IL-4, underscoring their biological significance as a primary source of this cytokine (63, 64). However, the *in vivo* effects of IL-4 on the immune response to tumors are complex, mainly due to the diversity of cellular targets influenced by IL-4. For instance, IL-4 exhibits a dual role in tumor immunity. Genetic overexpression of IL-4 in tumor cells has been shown to lead to tumor rejection (65–67). Conversely, IL-4 has also been found to promote tumor progression, either directly by facilitating the growth of tumor cells that express the IL-4 receptor or indirectly by triggering an immunosuppressive TME (68, 69). In our study, we provide compelling data showing that IL-3 robustly stimulates melanoma-associated basophils to produce elevated levels of IL-4. Importantly, our research also elucidates how basophil-derived IL-4 directly affects CD8^+^ T cells by remodeling their metabolism, boosting their effector functions, and prolonging their survival. This modulation of CTLs ultimately leads to an enhanced anti-tumor response. Our data underscore the critical role of basophil-derived IL-4 in enhancing CTL anti-tumor activity, shedding light on the intricate interplay between basophils and anti-tumor immunity. While the overall impact of basophil-derived IL-4 on tumor responses may be influenced by diverse factors, such as the composition of the TME, the type of cancer, and the stage of disease, the significance of high IL-4 expression in the TME has opened up new opportunities for therapeutic intervention. These findings hold great promise for the development of novel immunotherapeutic strategies to effectively combat cancer.

Stemming from this study, numerous opportunities emerge for harnessing T cell-basophil crosstalk mediated by IL-3 in the realm of cancer treatment. Adoptive T-cell therapy stands out as a highly favorable strategy, enabling the amplification of tumor-specific CD8^+^ T cells in the presence of IL-4, thereby acquiring superior anti-tumor immunity. Moreover, our investigations into intraperitoneal administration of recombinant IL-3, adoptive transfer of T cells overexpressing IL-3, and infusion of IL-3-activated basophils consistently demonstrate enhancement of anti-tumor activity. To further develop improved cancer therapies, we suggest considering IL-3 augmentation in the genetic modification of tumor-reactive CTLs, such as chimeric antigen receptor (CAR) T-cell therapy. Additionally, the integration of basophils or IL-4 into the activation and expansion processes of CAR-T cells prior to adoptive transfer represents an innovative direction. Collectively, our diverse findings significantly enhance our understanding of CTL-mediated anti-tumor mechanisms, unveiling new targets that hold promise for the development of pioneering cancer treatments.

## MATERIALS AND METHODS

### Mice

C57BL/6J (B6, Strain #000664), B6.129S7-Rag1tm1Mom/J (Rag1 KO, Strain #002216), C57BL/6-Tg(TcraTcrb)1100Mjb/J (OT1, Strain #003831), B6.PL-Thy1a/CyJ (Thy1.1, Strain #000406), and C57BL/6-Il4^tm1Nnt^/J (IL-4 knockout, Strain #002518) mice were obtained from The Jackson Laboratory. Thy1.1 OT1 mice were generated by crossing Thy1.1 mice with OT1 mice. Animals were maintained in specific-pathogen free animal facility at 22°C and 40–50% humidity, with a daylight cycle from 6 a.m. to 6 p.m. and were provided 6% fat food (LabDiet® 5K52) and water *ad libitum*. Animal experiments were conducted in accordance with procedures approved by the Institutional Animal Care and Use Committee of The Jackson Laboratory or the Animal Care and Use Committee of Shandong University. Age-matched male mice (approximately 8-10 weeks old) were randomly assigned to experimental groups for all experiments.

### Cell lines

The B16-OVA cell line was authenticated by whole genome and transcriptome sequencing and was kindly provided by Dr. Hildegund Ertl, with permission from its originators, Drs. Edith Lord and John Frelinger. B16-OVA cells were transduced with lentivirus expressing IL-3 (B16-IL3) or with empty vector only (B16-Mock), and puromycin was used to select for stably transduced tumor cells. The 293T cell line was originally obtained from the American Type Culture Collection (ATCC). The B16-OVA, B16-IL3, and B16-Mock cells were grown in RPMI-1640 medium containing 100 units/mL penicillin and 100 μg/mL streptomycin, supplemented with 10% fetal bovine serum (FBS). The 293T cells were maintained in Dulbecco’s Modified Eagle Medium (DMEM) containing 100 units/mL penicillin and 100 μg/mL streptomycin sulfate, as well as 10% FBS. All cells were cultured at 37°C in a humidified incubator in the presence of 5% CO_2_. All the cell lines were routinely monitored for mycoplasma and other microbial contamination.

### Tumor models

For the generation of melanoma-bearing mice, tumor cells (B16-OVA, B16-IL3 and B16-Mock) in 50 μl of phosphate-buffered saline (PBS) were injected subcutaneously (s.c.) into the right flank of mice (5×10^5^ cells/mouse), followed by treatments as indicated in figure legends. Tumor size was measured via caliper every 2 days starting from day 3 or 5 (as indicated), with double-blinding employed for tumor size assessment in mice. Tumor volume was calculated using the formula: V = π×W^2^×L/6, where width is the minor tumor axis (W) and length is the major tumor axis (L). The humane endpoint (euthanasia required) for a single tumor is volume >2,000 mm^3^. 2.5×10^5^ B16-IL3 or B16-Mock cells were intravenously (i.v.) injected into mice to induce the development of metastases in the liver.

### Tumor dissociation

Tumors were excised from mice at indicated days after tumor inoculation. Isolated tumor tissues were manually dissociated and subsequently incubated with 200 units/ml collagenase type I and 100 µg/ml DNase I in RPMI-1640 containing 10% FBS for 1h at 37°C. Single cell suspensions were obtained by passing digested tumors through a 70 μm cell strainer.

### Cell culture

Naive T cells from spleens and lymph nodes were purified by MACS microbeads (Miltenyi Biotec) and seeded in RPMI-1640 containing 10% FBS, 100 units/mL penicillin, 100 μg/mL streptomycin, 55 μM β-mercaptoethanol, 1% L-glutamine and 1 mM sodium pyruvate. They were cultured at 37°C in humidified incubators containing 5% CO_2_ and stimulated with 0.5 μg/mL αCD28 and 100 units/mL IL-2 on 5 μg/mL αCD3 pre-coated plates for 3 days. For treatment with IL-3 or IL-4, activated T cells were incubated with the indicated cytokines from day 1 to day 2 post stimulation. Activated T cells were subjected to flow cytometric analysis for cell markers, cytokine production, and survival. For low glucose treatment, glucose-free RPMI-1640 supplemented with either 11 mM (control) or 0.5 mM (low glucose) D-glucose was used. For extended stimulation, T cells were cultured with 0.5 μg/ml αCD28 and 100 units/mL IL-2 on 5 μg/mL αCD3 coated plates for an additional 3 days.

### Intracellular staining and flow cytometry

Intracellular staining was carried out as previously described (70). Briefly, T cells were incubated in complete medium supplemented with 1.0 μL/mL GolgiStop in the presence or absence of PMA and ionomycin for 4 hours. Then cells were fixed with BD Cytofix/Cytoperm Plus Fixation/Permeabilization Solution Kit at 4°C according to the manufacturer’s instructions. Flow cytometry staining was performed with fluorochrome-conjugated antibodies in PBS containing 1% FBS and 2 mM EDTA (surface stain) or Cytoperm/Wash (intracellular stain) on ice. Dead cells were excluded from analysis with either propidium iodide or LIVE/DEAD™ Fixable Aqua Dead Cell Stain Kit. Data were acquired on FACSymphony A5 flow cytometers (BD Biosciences) with FACSDiva software (v. 6.1.3) and analyzed by FlowJo software (v. 10.8.1).

### Quantitative RT-PCR

Total RNA was extracted from cells using the PureLink RNA Mini Kit (Invitrogen) and first-strand cDNAs from equivalent amounts of RNA for each sample were synthesized using the High-Capacity cDNA Reverse Transcription Kit using manufacturers’ recommendations. Quantitative RT-PCR was performed using PowerUp™ SYBR™ Green Master Mix (Applied Biosystems) in duplicate using the following primers: *Il3*-F: AGCTCCCAGAACCTGAACTCAAA; *Il3*-R: CAGAGTCATTCGCAGATGTAGGC; *Il4*-F: TTTGAACGAGGTCACAGGAGAA; *Il4*-R: CACCTTGGAAGCCCTACAGACG; *Actb*-F: AGAGGGAAATCGTGCGTGAC; *Actb*-R: CAATAGTGACCTGGCCGT.

### Retroviral transduction

Naive OT1 T cells (isolated from C57BL/6J OT1 mice) were stimulated with 0.5 μg/mL αCD28 and 100 units/mL IL-2 in 5 μg/mL αCD3 pre-coated plates. On day 1 of activation, cells were transduced with GFP reporting retroviruses expressing IL-3 or Mock control plasmids through centrifugal infection (2,500 rpm, 90 min) in the presence of 8 μg/mL polybrene and 20 mM HEPES, followed by incubation with the viruses overnight. 2 days post-transduction, IL-3- or Mock-expressing OT1 cells were sorted for GFP expression by fluorescence activated cell sorting (FACS) sorting on a FACSAria II sorter, and adoptively transferred into B16-OVA-bearing mice (5 days post-tumor inoculation (p.t.i)).

### Cellular therapy

For adoptive transfer of T cells, OT-1 or Thy1.1 OT1 cells were purified via CD8a (Ly-2) MACS microbeads (Miltenyi Biotec), unless otherwise indicated, and stimulated with αCD3 and αCD28 for one day. Cells were cultured for 2 additional days in the presence of treatments as indicated in figure legends. At day 3 following *in vitro* activation, the OT1 cells were washed with PBS and adoptively transferred into B16-OVA-bearing mice by i.v. injection at day 5 p.t.i (2×10^6^ cells/mouse). After T cell transfer, tumor size was measured every other day.

For therapy by basophils, 5×10^5^ B16-IL3 cells were subcutaneously implanted into Rag1 KO mice. On day 12 post inoculation, basophils were isolated from spleens and lymph nodes of the B16-IL3-melanoma-bearing Rag1 KO mice by FACS sorting. The isolated basophils were washed with PBS, and adoptively transferred into B16-OVA-bearing C57BL/6J mice on day 3 p.t.i., with a dosage of 2×10^6^ basophils per mouse). After basophil transfer, tumor size (measured every other day) and survival of recipient mice were recorded.

### Treatment of tumor-bearing mice

For IL-3 treatment conditions, IL-3 in 200 μl of PBS was injected into the peritoneal cavity of recipient mice every 2 days starting from day 3 p.t.i (300 ng/mouse). For depletion of CD8^+^ T cells, mice were intraperitoneally (i.p.) treated with neutralizing antibody against mouse CD8a (Leinco Technologies) or Rat IgG2a κ isotype control (100 μg/mouse) every 4 days beginning 5 days prior to tumor inoculation until the endpoint. For depletion of basophils, mice were i.p. treated with neutralizing antibody against mouse αFcεR1 (MAR-1, Life Technologies) or Armenian Hamster IgG isotype control (5 μg/mouse) at - 3, -2, -1, or 0 days prior to tumor injection and days 2, 5, and 10 p.t.i.

### Glucose uptake and metabolism assay

For glucose uptake assay *in vitro*, cells were incubated in complete medium with 5 μg/mL 2-NBDG at room temperature for 15 minutes and then analyzed by flow cytometry. Extracellular acidification rates (ECAR) and oxygen consumption rates (OCR) were determined by an XFe96 Extracellular Flux Analyzer (Seahorse Bioscience). Cells were plated in nonbuffered RPMI-1640 medium with 25mM glucose in Poly-D-Lysine pre-coated XFe96-well plates (2×10^5^ cells per well). Measurements were obtained under basal conditions.

### Survival of *in vitro* T cells

On day 3 of activation, CD8^+^ T cells were counted and equally replated in quadruplicate within 96-well plates, using RPMI-1640 media containing 10% FBS, 100 units/mL penicillin, 100 μg/mL streptomycin, 0.05 mM β-mercaptoethanol, 1% L-glutamine, and 1 μM sodium pyruvate. Propidium iodide staining was used to assess live cell number through flow cytometry analysis daily from day 3 to day 6. The cell survival rate was determined as the percentage of live cells relative to the number on day 3.

### *In vitro* cytotoxic assay

An assay for cellular *in vitro* cytotoxicity was conducted as previously described (13). Briefly, B16-OVA melanoma cells were pre-treated with 300 units/mL murine IFN-γ for 48 hours. Then, a single cell suspension was prepared. Target B16-OVA cells were generated by staining with 2 μM CFSE in PBS for 15 minutes at room temperature. *In vitro* activated CD8^+^ OT1 cells were co-cultured with 10,000 target cells per well in 96-well round bottom plates at the indicated T cell:tumor cell ratios for 12 hours at 37°C with 5% CO_2_. The cells were then trypsinized and 10,000 reference B16-OVA cells (labeled with 10 μM CFSE) were added, followed by staining with Po-ProTM-1 dead cell staining dye (Life Technologies). T cell killing efficiency was determined by flow cytometry and defined as the percentage of live tumor cells relative to reference cells.

### Transwell assay

Transwell chambers (Corning, 0.4 μm pore size) were placed in a 6-well plate. 3×10^5^ purified CD8^+^ T cells were simulated in complete medium supplemented with 0.5 μg/mL αCD28 and 100 units/mL IL-2 in the lower chamber that were pre-coated with 5 μg/mL αCD3, and 1.5×10^6^ bulk splenocytes were cultured in the same medium in the upper chamber. On day 3 post activation, CD8^+^ T cells in the lower chamber were collected and subjected to polyfunctionality analysis.

### Supernatant assay

Purified CD8^+^ T cells were cultured in complete medium supplemented with 0.5 μg/mL αCD28 and 100 units/mL IL-2 in 5 μg/mL αCD3 pre-coated plates for 2 days and then the conditioned supernatant was collected by centrifugation. Bulk splenocytes from Rag1 KO mice were cultured in complete medium supplemented with 100 units/mL IL-2 for 2 days and their supernatant was also collected by centrifugation. As indicated in figures, the conditioned supernatants from T cells or splenocytes plus half volume of fresh complete medium were used for the culture of CD8^+^ T cells or splenocytes from Rag1 KO mice. For antibody blockade experiments, indicated antibody (αIL-3, αIL-4, or isotype) was added into the conditioned supernatant to a final concentration of 5 μg/mL 2 hours prior to cell culture.

### RNA-sequencing and analysis

RNA-sequencing (RNA-seq) analysis was performed by The Jackson Laboratory Genome Technologies Scientific Service. Total RNA (3 independent samples per group) was isolated (QIAGEN miRNeasy mini extraction kit (QIAGEN) and cDNA was synthesized (High-Capacity cDNA Reverse Transcription Kit (Applied Biosystems)). RNA quality was assessed via RNA integrity (RIN) analysis (Bioanalyzer 2100, Agilent Technologies). Only RNA samples with a RIN value of 7 or higher were included in the analysis. RNA-seq libraries were generated from Poly(A) RNA capture using the Illumina TruSeq RNA Sample preparation kit v2. RNA-seq was performed in a 75-bp paired-end format with a minimum of 40 million reads per sample on the Illumina NextSeq 500 platform, according to the manufacturer’s instructions. RNA-seq reads were filtered and trimmed for quality scores >30 using a custom python script. The filtered reads were aligned to *Mus musculus* reference genome GRCm38 using RSEM (v1.2.12) (71) equipped with Bowtie2 (v2.2.0) (72) (command: rsem-calculate-expression -p 12 --phred33-quals --seed-length 25 --forward-prob 0 --time --output-genome-bam -- bowtie2). RSEM calculates expected counts and transcript per million (TPM). The expected counts from RSEM were used in the Bioconductor edgeR 3.20.9 package (73) to determine differentially expressed genes. Kyoto Encyclopedia of Genes and Genomes (KEGG) pathway enrichment analysis was performed using NetworkAnalyst (74, 75). Determination of the significantly different gene groups was performed with Gene Set Enrichment Analysis (GSEA) (76, 77). The following gene sets from the Molecular Signatures Database were used: systematic names, M9809 (Cytokine-Cytokine Receptor Reaction), M10617 (Inflammatory Response), M10267 (T-cell Activation), M17715 ( T-cell Proliferation), M17411 (JAK-STAT Signaling), M39736 (PI3K-AKT Signaling), M5562 (Tyrosine Phosphorylation of STAT Protein), and M11062 (Intracellular Signaling Transduction). Groups were deemed significant at an FDR *q*-value of 0.05 or below. Genes of interest for each group were extracted based on the core enrichment value provided by GSEA. RNA-seq results were further analyzed using Ingenuity Pathway Analysis (IPA, QIAGEN, version 01-22-01) for Upstream Regulator Analysis of differentiation expression genes identified in this study. Bonferroni method was used to correct the detection *p*-value and set 0.05 as the significant threshold.

### Tumor patient survival analysis

Kaplan–Meier (KM) analyses were performed to study the association between tumor patient survival and expression of basophil markers genes in melanoma using the KM Plotter online tool (78) (http://kmplot.com/analysis/). Gene expression analyses of tumor-resident basophils (by CD123, CCR3 and FcεR1) were performed. Patients’ overall survival and progression free survival were analyzed up to 60 months. Based on combined gene expression levels of selected basophil markers, patients were grouped into the top tertile (T3) and lower tertile (T1), while patients in the middle tertile (T2) were excluded. Markers (Gene Probe ID): CD123 (206148_at); CCR3 (208304_at); FcεR1 (211734_s_at).

### Statistical analysis

Sample sizes with sufficient statistical power for all studies were determined through preliminary pilot experiments. Data collected from flow cytometry was analyzed using FlowJo v10 on appropriate gated cells after exclusion of doublets and dead cells. Statistical analysis was performed using Graphpad Prism v8 and represented as mean ± standard error of mean (SEM). Statistical methods can be found in figure legends, as well as samples size and number of independent replicates. Briefly, comparisons between two groups were determined using unpaired two-tailed Student’s *t*-test. When multiple groups were compared, one-way ANOVA was applied. Two-way ANOVA was used in tumor growth analysis. For survival analysis, *p* values were calculated using the Log-rank test. Correlation analysis was performed using simple linear regression. The level of significance was set to *p* < 0.05. *p* ≥ 0.05 was considered no significance (ns), and *p* < 0.05 was considered significant. \**p* < 0.05, \*\**p* < 0.01, \*\*\**p* < 0.001.

## Author Contributions

JW and C-HC designed the experiments. JW, CLM, XL, FH, JJW, and JDS performed all the experiments. JW, CLM, XL, FH, TK, JJW, NAL, DC, and C-HC analyzed all the data. All authors contributed to writing the manuscript.

## Acknowledgements

We thank Will Schott, Danielle Littlefield, and Krystal-Leigh Brown for procedural expertise with flow cytometry experiments. We gratefully acknowledge the contribution of Flow Cytometry, Computational Sciences, and Genome Technologies Scientific Services at The Jackson Laboratory. Drs. Edison Liu, Karolina Palucka, Guangwen Ren, and David Serreze for reagents and advice. Artwork is created with BioRender (BioRender.com) and Figdraw (https://www.figdraw.com/static/index.html). We are also extremely grateful for the funding support from National Institutes of Health Grant (CA034196), American Cancer Society Research Grant (IRG-16-191-33), The Jackson Laboratory Director’s Innovation Fund (19000-21-07), The V Foundation (V2021-036), the AAI Careers in Immunology Fellowship Program, the Pyewacket Fund, the National Natural Science Foundation of China (82371753), the Shandong Provincial Science Fund for Excellent Young Scientists Fund Program (2023HWYQ-036), and the Shandong Provincial Natural Science Foundation (ZR2023MH163).

## Supplemental Figure Legend

**Figure S1. Tumor-infiltrating T lymphocytes progressively become dysfunctional during tumor growth**

(**A-F**) Relative mean fluorescence intensity (MFI) of PD-1 (**A**), TIM-3 (**B**), T-bet (**C**), IFN-γ (**D**), TNF-α (**E**), and IL-2 (**F**) in CTLs within B16-OVA melanoma at day 8, 13, 18, and 23 p.t.i were analyzed by flow cytometry. The MFI was normalized to the corresponding average value of 8-day p.t.i (n = 4-6). Statistical data are presented as the mean ± SEM, and significance was determined by one-way ANOVA. ns, no significance; \*\**p* < 0.01; and \*\*\**p* < 0.001.

**Figure S2. IL-3 production in activated effector T cells**

(**A, B**) Representative flow cytometry plots depicting IL-3 production by naive and *in vitro* activated CD8^+^ (**A**) and CD4^+^ (**B**) T cells.

(**C, D**) IL-3 MFI of CD8^+^ (**C**) and CD4^+^ (**D**) T cells in B16-OVA melanoma at day 8, 11, 14, and 17 p.t.i was analyzed by flow cytometry. IL-3 MFI is presented as the mean ± SEM (n = 7).

(**E**) IL-3 MFI in day 3- and day 6-activated CD8^+^ T cells with anti-CD3/CD28 stimulation. The data was normalized to the average value of Day 3 (n = 3).

(**F**) Relative IL-3 MFI of *in vitro* activated CD8^+^ T cells under glucose restriction. Statistical data are presented as the mean ± SEM.

(**G**) Relative mRNA level of *Il3* gene in *in vitro* activated CD8^+^ T cells under glucose restriction was analyzed by RT-PCR. Statistical data are presented as the mean ± SEM.

Statistical significance was determined by one-way ANOVA for **C** and **D**; and Student’s *t*-test for **E**-**G**. ns, no significance; \**p* < 0.05, \*\**p* < 0.01, and \*\*\**p* < 0.001.

**Figure S3. IL-3 does not directly affect metabolism or effector function of CD8^+^ T cells**

(**A**) Glucose uptake of *in vitro* CD8^+^ T cells treated with PBS or IL-3 was monitored by 2-NBDG. (**B-D**) Measurements of basal ECAR (**B**) and OCR (**C**) of *in vitro* CD8^+^ T cells treated with PBS or IL-3. ECAR/OCR ratio (**D**) was also determined (n = 10).

(**E-N**) Purified CD8^+^ T cells were activated with anti-CD3/CD28 in the presence of PBS or IL-3 for 3 days. Frequencies or MFI measurements of CD44 (**E**), PD-1 (**F**), TIM-3 (**G**), CD25 (**H**), T-bet (**I**), Ki67 (**J**), TNF-α (**K**), IL-2 (**L**), IFN-γ (**M**), and GZMB (**N**) of the T cells were examined following activation (n = 3).

(**O**) Survival of *in vitro* activated CD8^+^ T cells following 3 days-treatment with PBS or IL-3 (n = 3). Statistical data are presented as the mean ± SEM, and significance was determined by Student’s *t*-test (**B-N**) or two-way ANOVA (**O**). ns, no significance; \**p* < 0.05.

**Figure S4. Co-culture with bulk splenocytes potentiates anti-tumor activity of CD8^+^ T cells**

(**A**) Purified CD8^+^ T cells (pCD8) and CD8^+^ T cells with bulk splenocytes (bCD8) were activated with anti-CD3/CD28 for 3 days. Glucose uptake was monitored by 2-NBDG. (**B-D**) Measurements of basal ECAR (**B**) and OCR (**C**) of cells. ECAR/OCR ratio (**D**) was also determined (n = 10).

(**E**) IFN-γ production by pCD8 and bCD8 cells was examined. Representative flow cytometry plots of IL-3 are presented.

(**F**) Survival of pCD8 and bCD8 cells following 3 days-activation (n = 3).

(**G, H**) Activated pCD8 and bCD8 Thy1.1 OT1 cells were adoptively transferred into B16-OVA-melanoma-bearing mice at day 10 p.t.i (**G**). Frequency of the transferred T cells in tumors were evaluated 2 and 4 days later (**H**) (n = 5-7).

(**I**) *In vitro* cytotoxicity of pCD8 and bCD8 OT1 cells. B16-OVA cells were co-cultured with activated pCD8 or bCD8 OT1 cells at the indicated ratios for 12 hours and survival of the tumor cells was examined (n = 3).

(**J, K**) Activated pCD8 and bCD8 OT1 cells were adoptively transferred into mice with B16-OVA-melanoma at day 5 p.t.i and the tumor size was recorded (**J**). Tumor growth curves (**K**) are presented (n = 6-12 mice/group).

(**L, M**) Purified CD8^+^ T cells were activated and cultured with conditioned medium of Rag1 KO mouse splenocytes treated with indicated PBS, IL-3, IL-5, or GM-CSF. IFN-γ production (**L**) and survival (**M**) of the cells were evaluated (n = 3).

Statistical significance was determined by Student’s *t*-test (**B**-**D**) or two-way ANOVA (**F**, **H**, **I**, **K**, and **M**). ns, no significance; \**p* < 0.05, \*\*\**p* < 0.001.

**Figure S5. Identification of the cellular targets of IL-3**

(**A**) The gating strategy for isolation of cell populations from the spleen of Rag1 KO mice by FACS.

(**B**) The purified cell subsets were cultured in the presence of IL-3 for 2 days and then the conditioned supernatant was collected for culture of CD8^+^ T cells. IFN-γ production by the CD8^+^ T cells was examined. Representative flow cytometry plots are presented.

(**C**) The gating strategy for isolation of basophils from Rag1 KO mice by FACS.

**Figure S6. The effects of IL-4 on genes expression and activity of CD8^+^ T cells**

(**A, B**) RNA-seq analysis of activated CD8^+^ T cells following incubation with conditioned supernatant from culture of basophils with IL-3 *versus* PBS. Kyoto Encyclopedia of Genes and Genomes (KEGG) pathway analysis was performed. Volcano plots for differently expressed genes (**A**) and the top10 regulated pathways (**B**).

(**C-F**) GSEA plots of activated CD8^+^ T cells following incubation with conditioned supernatant from culture of basophils with IL-3 *versus* PBS (n = 3 mice per group).

Enrichment of gene sets in T-cell Activation (**C**), T-cell Proliferation (**D**), Tyrosine Phosphorylation of STAT Protein (**E**), and Intracellular Signaling Transduction (**F**).

(**G**) Glucose uptake of *in vitro* CD8^+^ T cells treated with PBS or IL-4 was monitored by 2-NBDG. (**H-J**) Measurements of basal ECAR (**H**) and OCR (**I**) of *in vitro* CD8^+^ T cells treated with PBS or IL-4. ECAR/OCR ratio (**J**) was calculated (n = 10).

(**K-S**) Purified CD8^+^ T cells were activated with anti-CD3/CD28 in the presence of PBS or IL-4. Frequency or MFI of CD44 (**K**), PD-1 (**L**), TIM-3 (**M**), CD25 (**N**), T-bet (**O**), Ki67 (**P**), TNF-α (**Q**),

IL-2 (**R**), and GZMB (**S**) expression was examined following activation (n = 3). Statistical data are presented as the mean ± SEM, and significance was determined by Student’s *t*-test. ns, no significance; \*\**p* < 0.01, and \*\*\**p* < 0.001.

**Supplemental Table 1.**
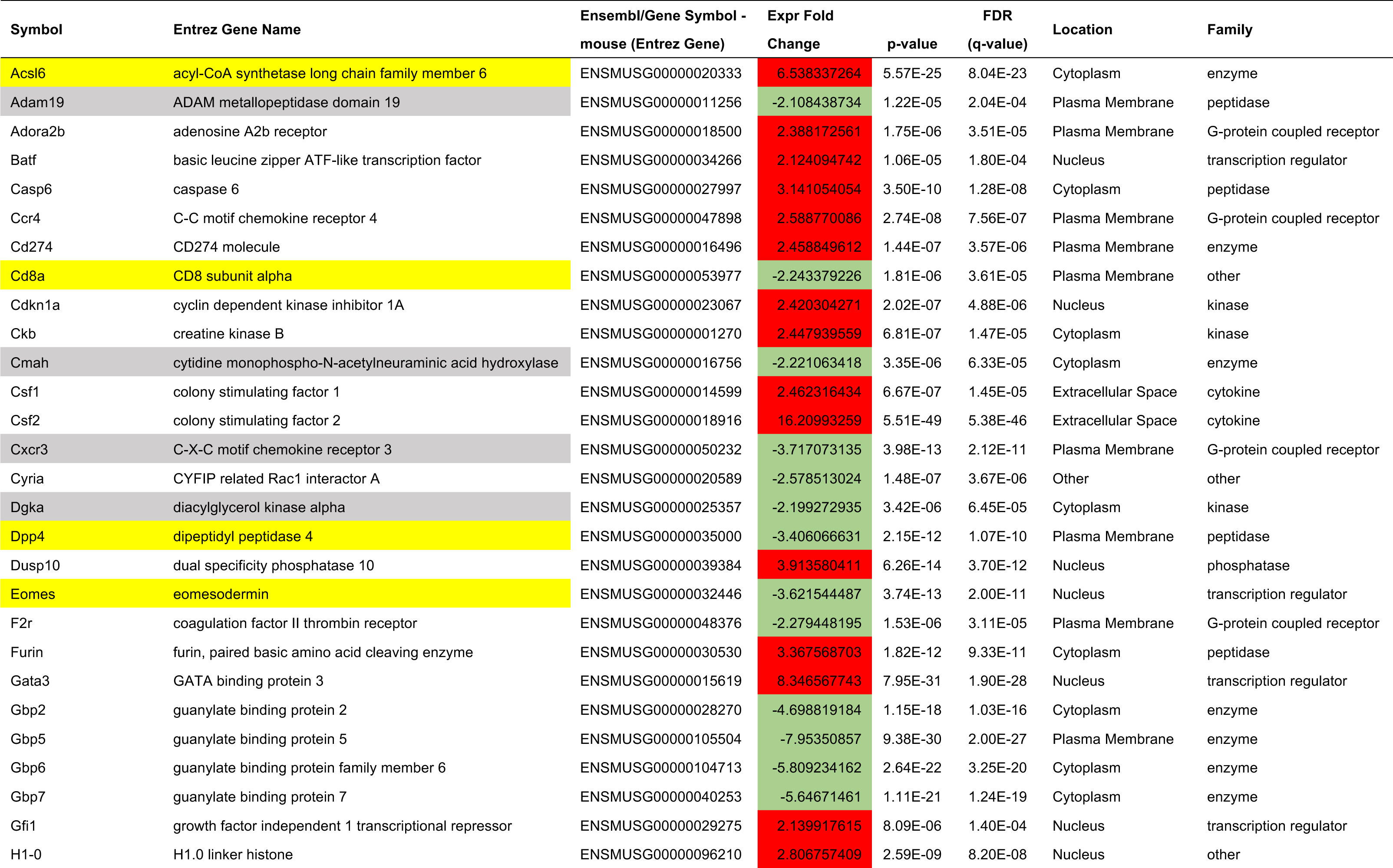
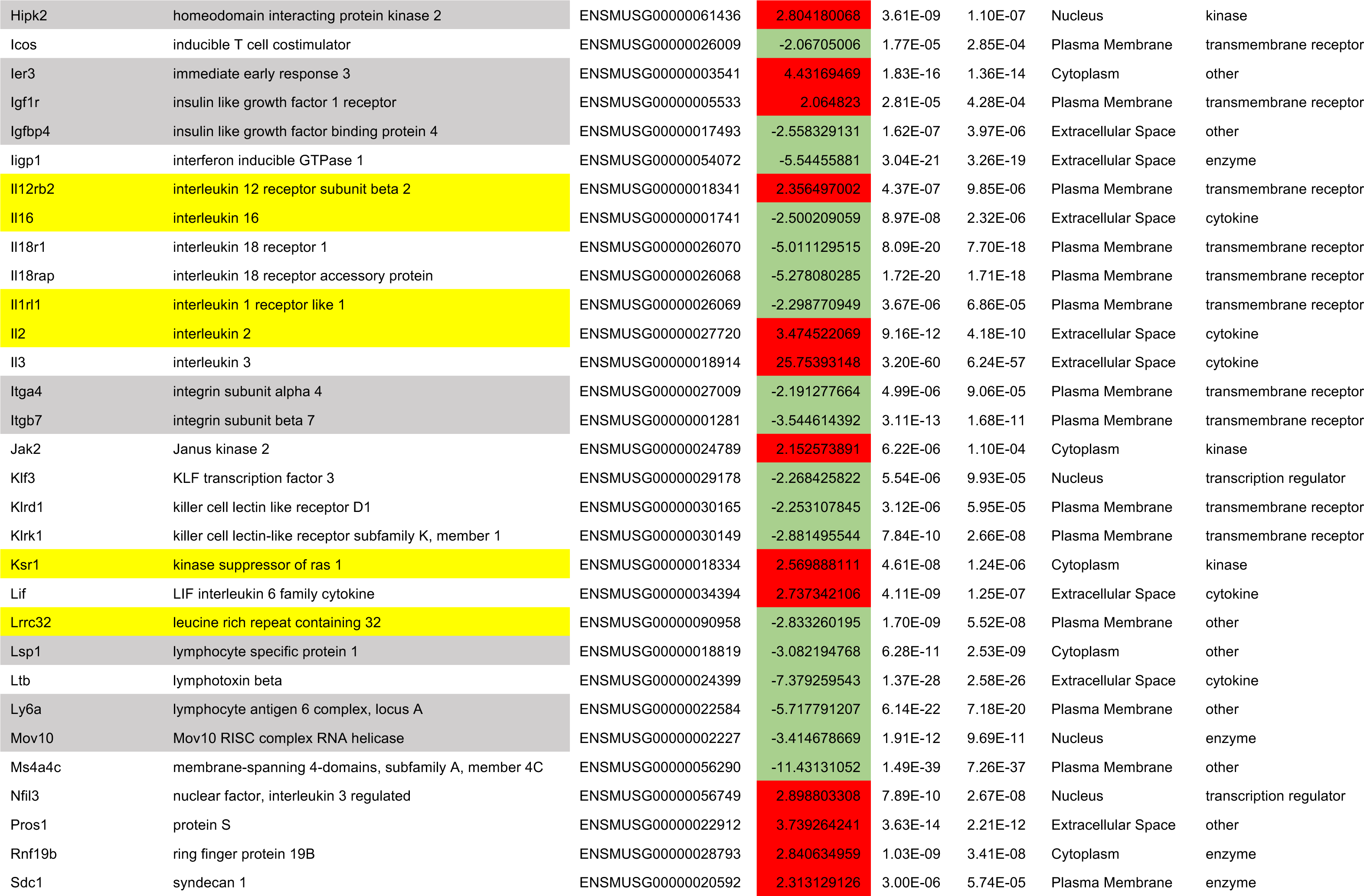
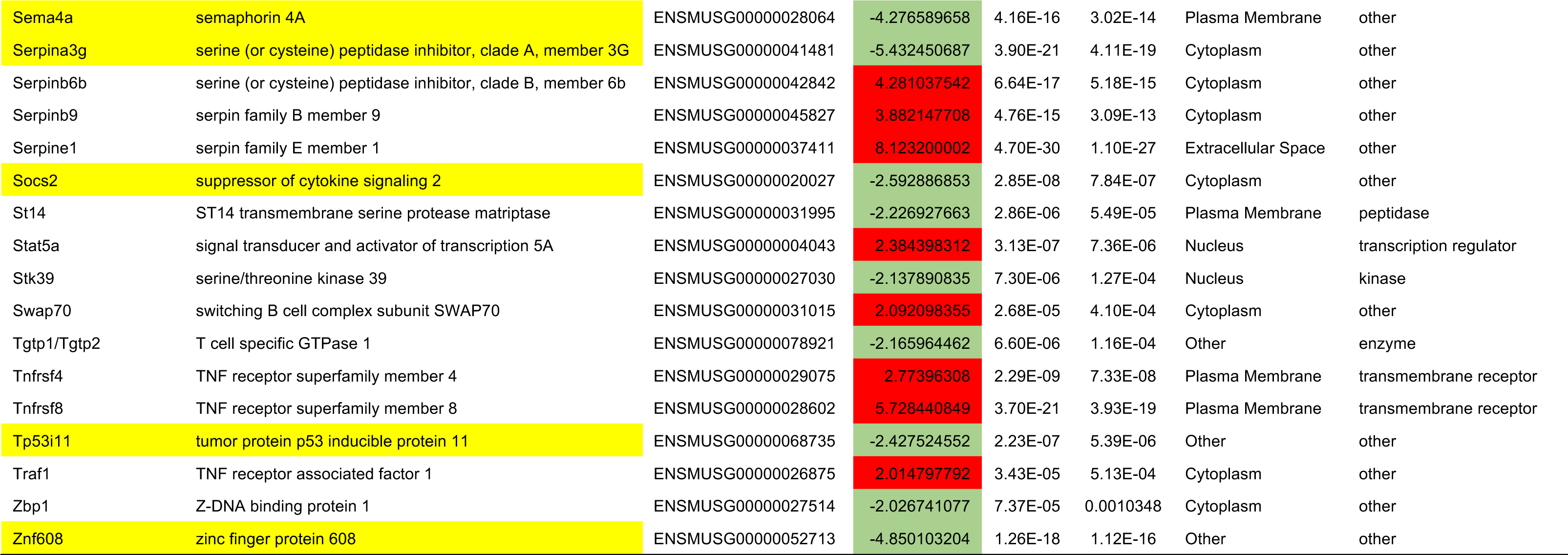
Impacted genes within the IL-4 regulatory network.

