## Supplemental figures 1-6 for "IL-3-driven T cell-basophil crosstalk enhances anti-tumor immunity"

### Figure S1

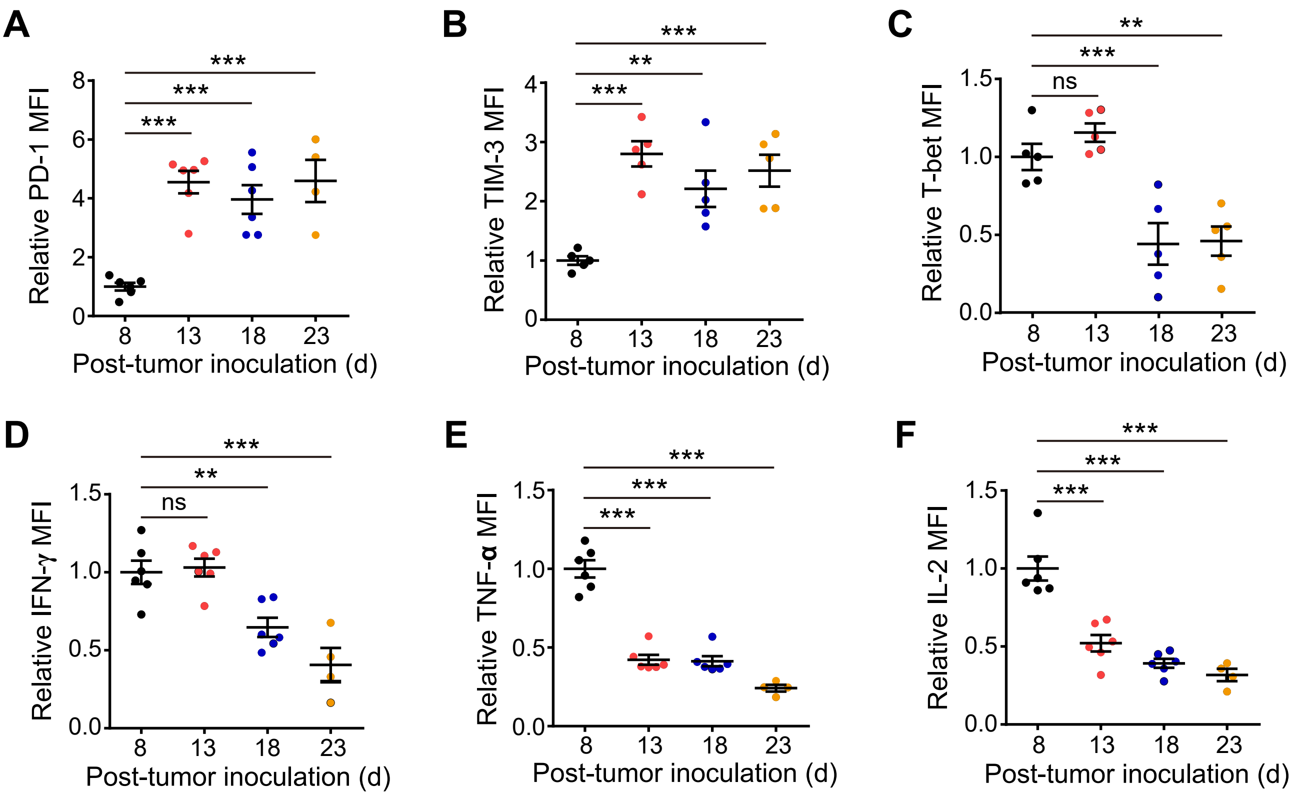
A
Relative PD-1 MFI
Post-tumor inoculation (d)
8
13
18
23
***
***
***
B
Relative TIM-3 MFI
Post-tumor inoculation (d)
***
**
***
C
Relative T-bet MFI
Post-tumor inoculation (d)
**
***
ns
D
Relative IFN-γ MFI
Post-tumor inoculation (d)
***
**
ns
E
Relative TNF-α MFI
Post-tumor inoculation (d)
***
***
***
F
Relative IL-2 MFI
Post-tumor inoculation (d)
***
***
***

### Figure S2

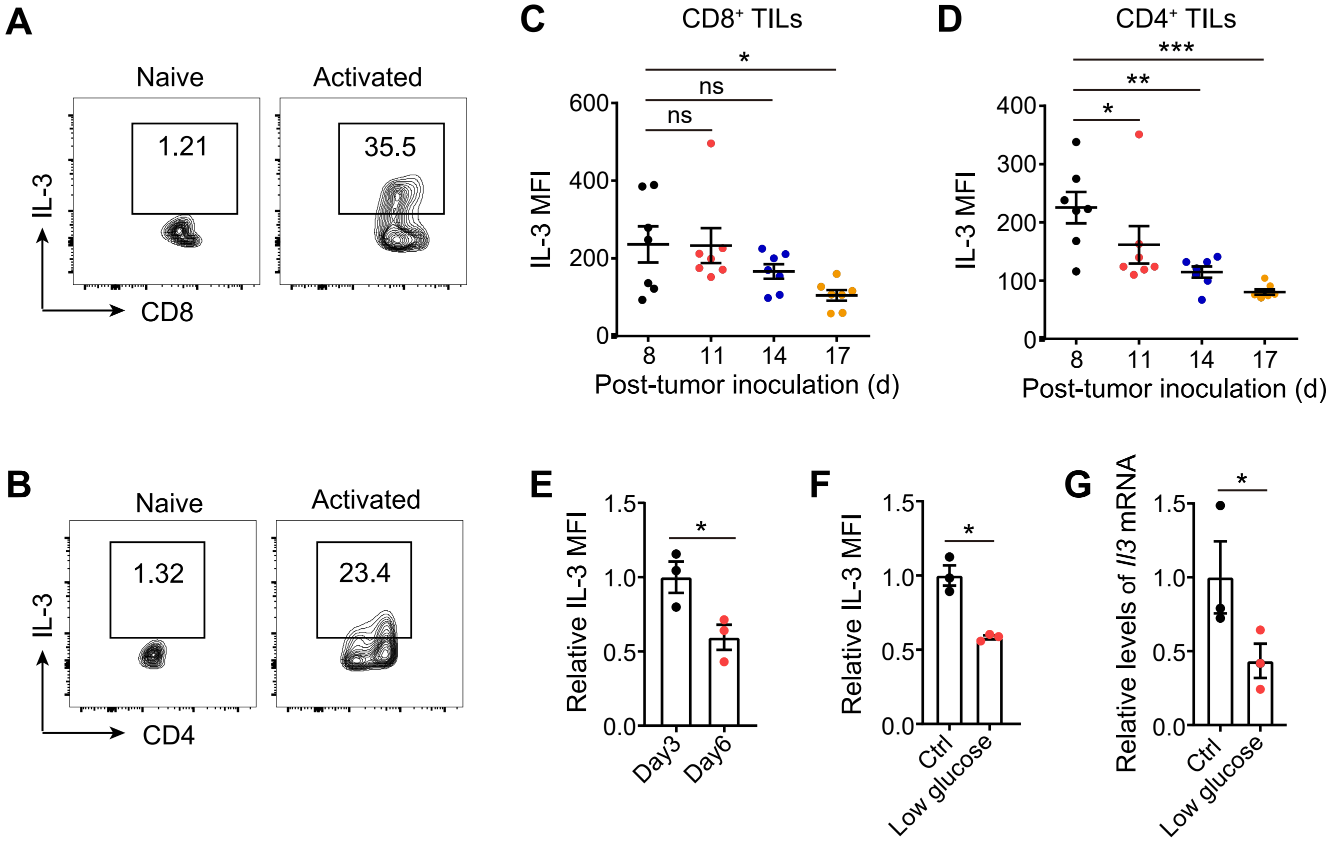
A
Naive
Activated
1.21
35.5
IL-3
CD8
B
Naive
Activated
1.32
23.4
IL-3
CD4
C
CD8⁺ TILs
IL-3 MFI
Post-tumor inoculation (d)
ns
ns
*
D
CD4⁺ TILs
IL-3 MFI
Post-tumor inoculation (d)
*
**
***
E
Relative IL-3 MFI
Day3
Day6
*
F
Relative IL-3 MFI
Ctrl
Low glucose
*
G
Relative levels of Il3 mRNA
Ctrl
Low glucose
*

### Figure S3

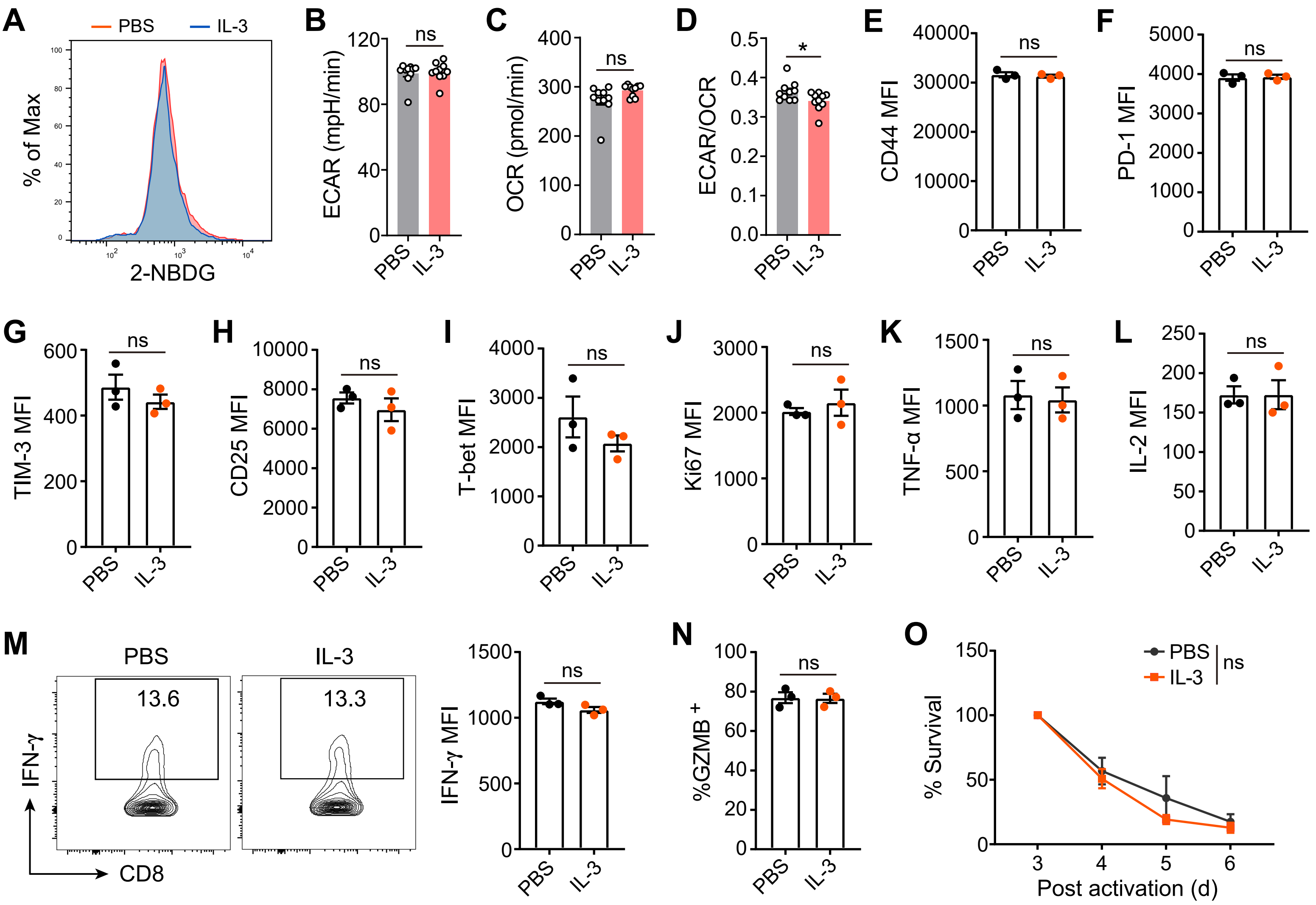
A
PBS
IL-3
% of Max
2-NBDG
B
ECAR (mpH/min)
ns
PBS
IL-3
C
OCR (pmol/min)
ns
D
ECAR/OCR
*
E
CD44 MFI
ns
F
PD-1 MFI
ns
G
TIM-3 MFI
ns
H
CD25 MFI
ns
I
T-bet MFI
ns
J
Ki67 MFI
ns
K
TNF-α MFI
ns
L
IL-2 MFI
ns
M
PBS
IL-3
13.6
13.3
IFN-γ
CD8
IFN-γ MFI
ns
N
%GZMB +
ns
O
PBS
IL-3
ns
% Survival
Post activation (d)

### Figure S4

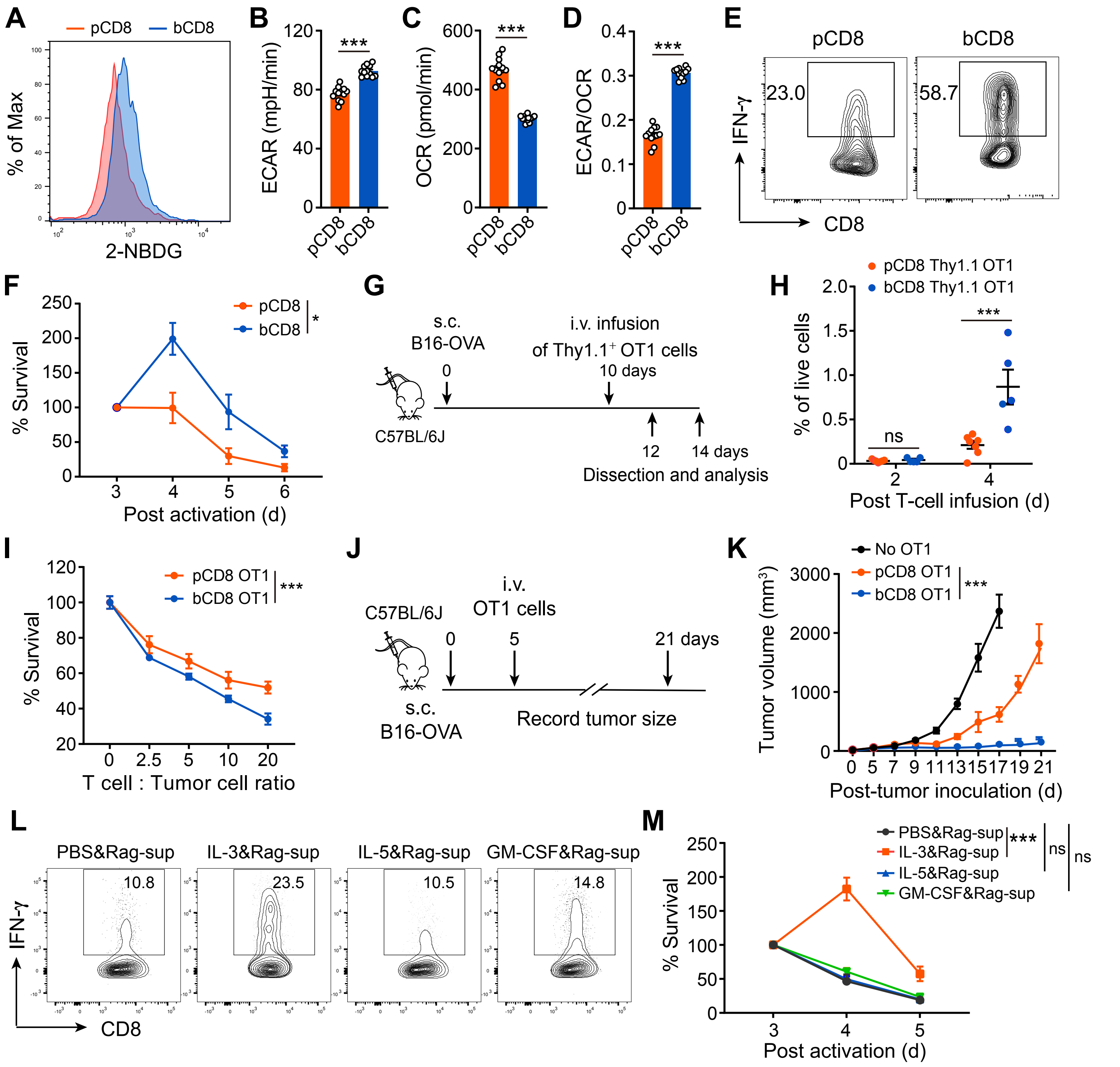
A
pCD8
bCD8
% of Max
2-NBDG
B
ECAR (mpH/min)
***
pCD8
bCD8
C
OCR (pmol/min)
***
D
ECAR/OCR
***
E
pCD8
bCD8
IFN-γ
CD8
23.0
58.7
F
% Survival
pCD8
bCD8
*
Post activation (d)
G
s.c.
B16-OVA
i.v. infusion
of Thy1.1+ OT1 cells
0
10 days
C57BL/6J
12
14 days
Dissection and analysis
H
pCD8 Thy1.1 OT1
bCD8 Thy1.1 OT1
% of live cells
ns
***
Post T-cell infusion (d)
I
% Survival
pCD8 OT1
bCD8 OT1
***
T cell : Tumor cell ratio
J
C57BL/6J
i.v.
OT1 cells
0
5
21 days
s.c.
B16-OVA
Record tumor size
K
No OT1
pCD8 OT1
bCD8 OT1
***
Tumor volume (mm3)
Post-tumor inoculation (d)
L
PBS&Rag-sup
IL-3&Rag-sup
IL-5&Rag-sup
GM-CSF&Rag-sup
10.8
23.5
10.5
14.8
IFN-γ
CD8
M
PBS&Rag-sup
IL-3&Rag-sup
IL-5&Rag-sup
GM-CSF&Rag-sup
***
ns
ns
% Survival
Post activation (d)

### Figure S5

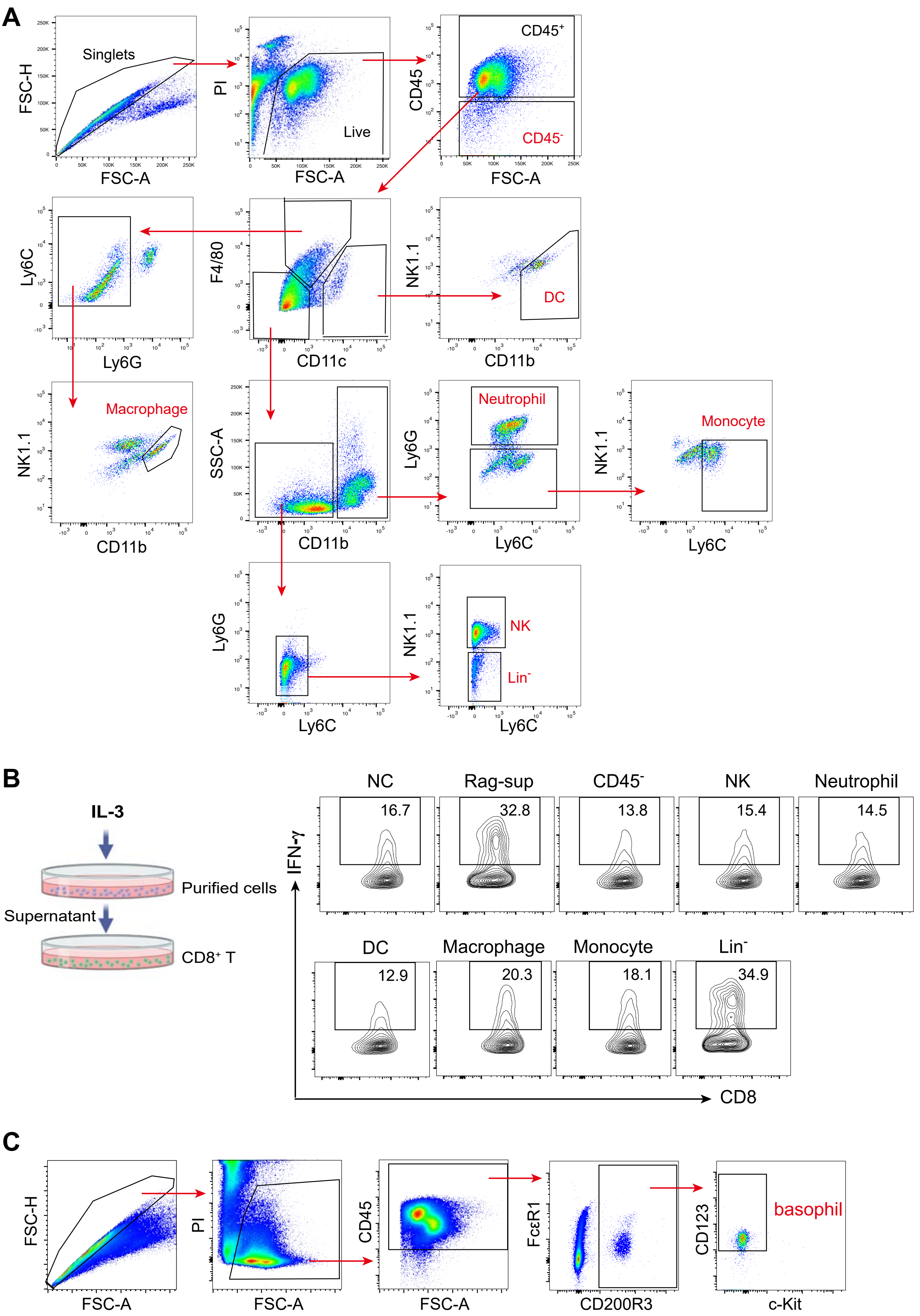
A
Singlets
FSC-H
FSC-A
PI
Live
CD45
CD45⁺
CD45⁻
Ly6C
Ly6G
F4/80
CD11c
NK1.1
CD11b
DC
Macrophage
SSC-A
Neutrophil
Monocyte
NK
Lin⁻
B
IL-3
Purified cells
Supernatant
CD8⁺ T
IFN-γ
CD8
NC
Rag-sup
CD45⁻
NK
Neutrophil
16.7
32.8
13.8
15.4
14.5
DC
Macrophage
Monocyte
Lin⁻
12.9
20.3
18.1
34.9
C
FSC-H
FSC-A
PI
CD45
FcεR1
CD200R3
CD123
c-Kit
basophil

### Figure S6

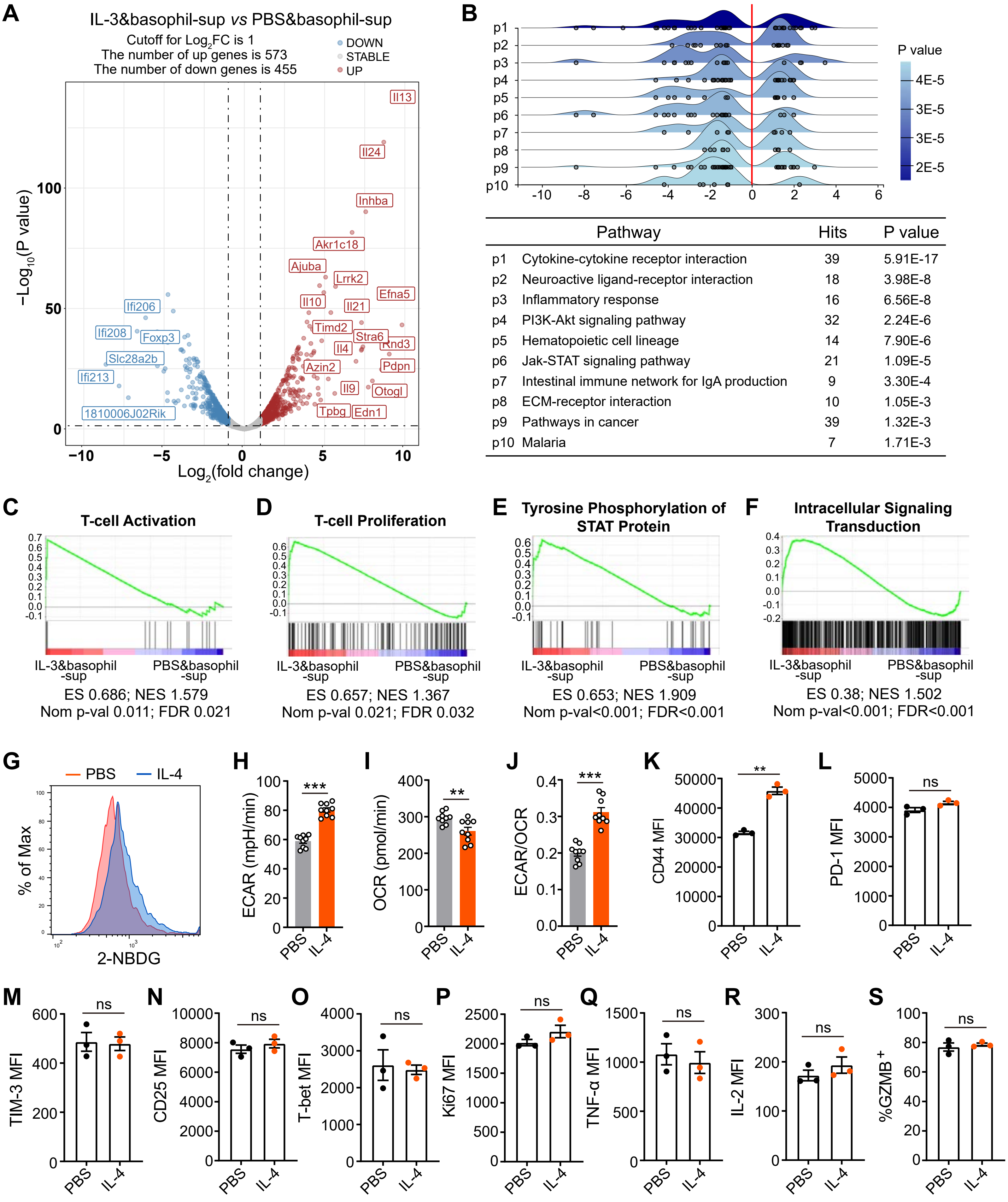


| | Pathway | Hits | P value |
|---|---|---|---|
| p1 | Cytokine-cytokine receptor interaction | 39 | 5.91E-17 |
| p2 | Neuroactive ligand-receptor interaction | 18 | 3.98E-8 |
| p3 | Inflammatory response | 16 | 6.56E-8 |
| p4 | PI3K-Akt signaling pathway | 32 | 2.24E-6 |
| p5 | Hematopoietic cell lineage | 14 | 7.90E-6 |
| p6 | Jak-STAT signaling pathway | 21 | 1.09E-5 |
| p7 | Intestinal immune network for IgA production | 9 | 3.30E-4 |
| p8 | ECM-receptor interaction | 10 | 1.05E-3 |
| p9 | Pathways in cancer | 39 | 1.32E-3 |
| p10 | Malaria | 7 | 1.71E-3 |
